# Genetic and functional diversity of β-*N*-acetylgalactosamine residue-targeting glycosidases expanded by deep-sea metagenome

**DOI:** 10.1101/2023.07.28.550916

**Authors:** Tomomi Sumida, Satoshi Hiraoka, Keiko Usui, Akihiro Ishiwata, Toru Sengoku, Keith A Stubbs, Katsunori Tanaka, Shigeru Deguchi, Shinya Fushinobu, Takuro Nunoura

**Author notes:** Corresponding authors: Tomomi Sumida, Shinya Fushinobu.

## Abstract

β-*N*-Acetylgalactosamine-containing glycans play essential roles in several biological processes, including cell adhesion, signal transduction, and immune responses. β-*N*-Acetylgalactosaminidases hydrolyze β-*N*-acetylgalactosamine linkages of various glycoconjugates. However, their biological significance remains ambiguous, primarily because only one type of enzyme, exo-β-*N*-acetylgalactosaminidases that specifically act on β-*N*-acetylgalactosamine residues, has been documented so far. In this study, we identified three novel glycoside hydrolase families distributed among all three domains of life and characterized eight novel β-*N*-acetylgalactosaminidases and β-*N*-acetylhexosaminidase through sequence-based screening of deep-sea metagenomes and subsequent searching of public protein databases. Despite low sequence similarity, the crystal structures of these enzymes demonstrate that all enzymes share a prototype structure and diversify their substrate specificities (endo-, dual-endo/exo-, and exo-) through the accumulation of mutations and insertional amino acid sequences. The diverse β-*N*-acetylgalactosaminidases reported in this study could facilitate the comprehension of their structures and functions and present novel evolutionary pathways for expanding their substrate specificity.

---

Beta-*N*-acetylgalactosamine (β-GalNAc)-containing glycans, such as glycoconjugates of polysaccharides (1), glycolipids (2, 3), *N*- and *O*-linked glycans (4, 5), O-antigen (6), and chondroitin sulfate (7), are ubiquitous and crucially contribute to various biological processes, including cell adhesion, signal transduction, cross-interactions with functional membrane components, formation of the cell envelope and maintenance of its stability, immunomodulation, and immune responses (1–7). The regulatory function of these glycans is attributed to their structural diversity, which differ in carbohydrate constituents (namely, glucose, galactose) and molecular architecture (α- or β-linkages, linear or branched) (1–8).

Beta-*N*-acetylgalactosaminidases (β-NGAs) hydrolyze the different β-GalNAc linkages of various glycans to modulate the length, combination, and abundance of glycans. This catalytic activity requires the β-NGAs to possess diverse substrate specificities, but only two enzymes, exo-β-NGA and exo-β-*N*-acetylhexosaminidase (exo-β-HEX), possessing distinctive substrate specificity and sequence, have been evidenced to hydrolyze β-GalNAc. Exo-β-NGAs have a strict substrate specificity for the non-reducing terminal β-GalNAc and are classified into the glycoside hydrolase (GH) family GH123 (9) of the Carbohydrate-Active Enzymes (CAZy) database (10, 11). As β-GalNAc is prevalent in various glycans across the three domains of life (Bacteria, Archaea, and Eukarya) and in different ecological niches (1–8), β-NGA activity is expected to follow a similar distribution pattern. However, these enzymes have been identified in only three bacterial species, specifically associated with microbe-host interactions in both terrestrial soil and human gut environments (namely, NgaP from *Paenibacillus* sp. TS12, CpNga123 from *Clostridium perfringens*, and BvGH123 from *Phocaeicola vulgatus*) (9, 12, 13), with no reported origin in archaea or eukaryotes, and endo-β-NGA has not been reported. Meanwhile, exo-β-HEXs hydrolyze the non-reducing terminal of β-GalNAc as well as β-GlcNAc, and are classified into the family GH20 (14–16). More than 100 exo-β-HEXs from all three domains of life have been functionally analyzed, with no report of endo-β-HEXs.

Given the limited extant knowledge on β-NGAs, it is imperative to further identify and functionally characterize novel β-NGAs. These endeavors are critical for comprehensively understanding the complex phenomena associated with β-GalNAc-mediated biological processes. Recently, culture-independent metagenomic exploration of novel glycosidases has substantially augmented our conception of carbohydrate-related enzymes. Function-based screening of diverse biological resources revealed several novel glycosidase families, including GH148 from volcanic soil (17), GH156 from a thermal hot spring (18), GH165 from agricultural soil (19), and GH173 and CBM89 from the capybara intestine (20). Moreover, sequence-based screening has enabled the analysis of much larger metagenomic sequencing datasets than functional screening, yielding more candidate sequences and facilitating the discovery of novel enzymes with distinct characteristics (21).

The deep-sea environments, characterized by unique features and distinct bacterial flora (22), have rarely been surveyed owing to the challenges associated with sampling from these regions. The deep-sea metagenome is a promising frontier for enzyme discovery (23, 24). Therefore, here we aimed to use a sequence-based screening approach of deep-sea microbial assemblages to explore the functional diversity of β-NGA activity. Our deep-sea sediment metagenomic (DSSM) analysis and domain search yielded three novel β-NGA gene families that are phylogenetically distinct from GH123 exo-β-NGAs. The biochemical and structural characterization of these enzymes not only unveiled their functional diversity but also shed light on their monophyletic evolutionary history, providing valuable insights into the mechanisms underlying β-GalNAc-mediated biological processes.

## Results

### Discovery of a novel β-NGA with both endo- and exo-glycosidase activities

Using four metagenomic datasets derived from microbial assemblages in deep-sea abyssal sediments and a domain-based search, we retrieved three candidate complete coding sequences (CDSs) (Gene ID: *dssm_1–3,* tentative Protein ID: DSSM_1–3), which exhibited low sequence identity (15–26%) to all known GH123 exo-β-NGA genes (*NgaP, CpNga123, and BvGH123*) (Extended Data Fig. 1a, Supplementary Data S1). Sequence alignments demonstrated that the consecutive catalytic “DE” motif of family GH123, comprising an aspartic acid (stabilizer of the 2-acetamido group of the substrate) and a glutamic acid (acid/base) (9), was present in the *dssm_2* and *dssm_3* sequences but not in *dssm_1* (Extended Data Fig. 1b, green box and asterisk).

AlphaFold2 (25) was used to predict the structure of the candidate proteins (Fig. 1a). DSSM_1 was structurally distinct from the GH123 exo-β-NGAs (Fig. 1a, b) and consisted of five domains, among which only domain 2 (β-sandwich) displayed a degree of structural similarity to the N-terminal domain of GH123 exo-β-NGAs. Domain 4 ((β/α)_8_ barrel) was similar to cycloisomaltooligosaccharide glucanotransferase (PDB, 3WNM). By contrast, the predicted structures of DSSM_2 and DSSM_3 were similar to those of GH123-β-NGAs, and they all shared DUF4091, a presumed domain region whose function remains undetermined (Fig. 1b, green, Extended Data Fig. 1b, underlined).

**Fig. 1.**
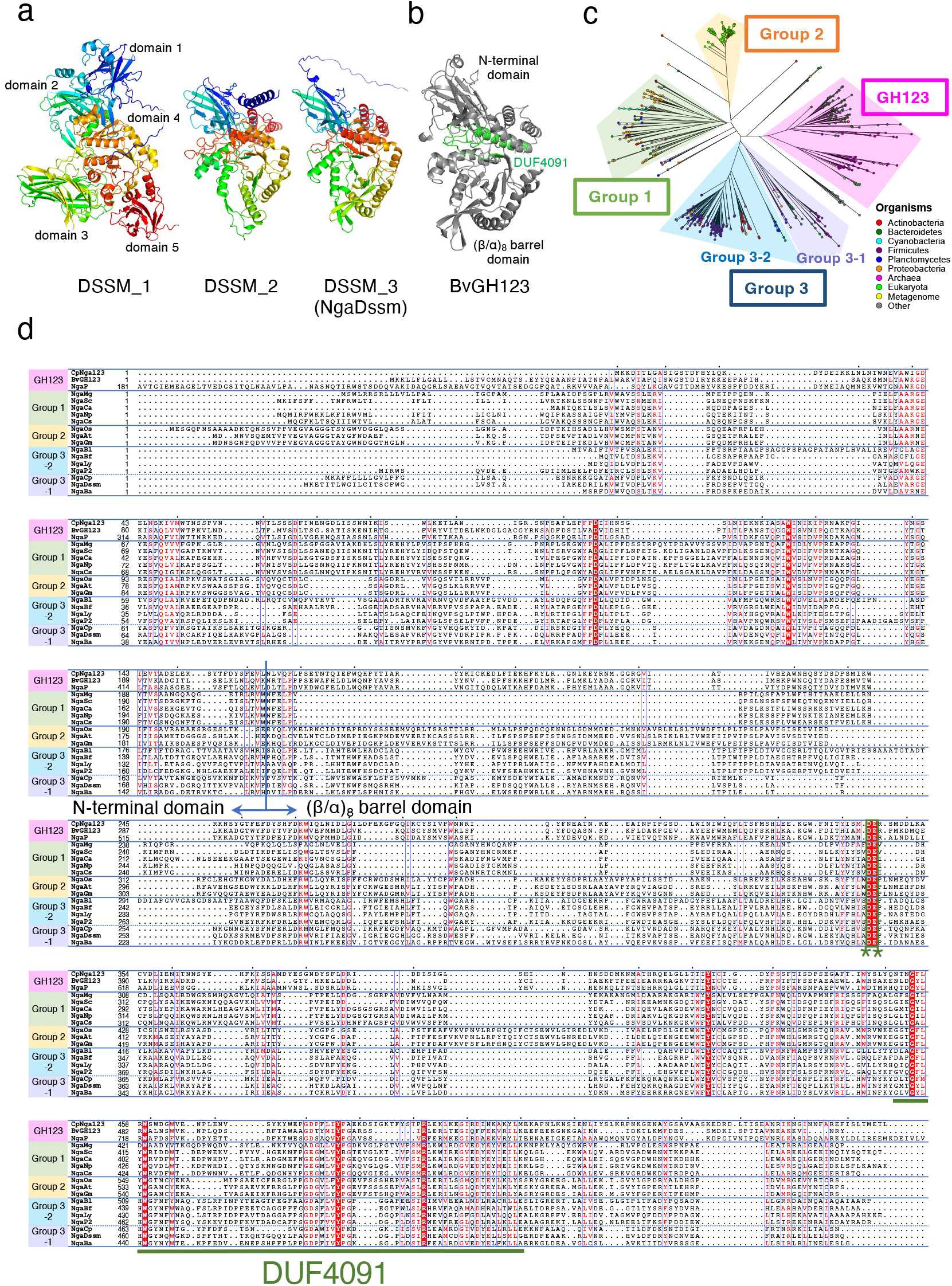
Candidate β-NGA sequences retrieved from deep-sea sediment metagenomes and phylogenetic and sequence diversities of β-NGA candidates. **a**, Overall structures of β-NGA candidates retrieved from deep-sea sediment metagenomes. The structures were predicted using AlphaFold2. **b**, The structure of BvGH123. DUF4091 is colored in green. **c**, Phylogenetic tree of β-NGA candidates retrieved from deep-sea sediment metagenomes and the Pfam database. **d**, Alignment of the β-NGA gene sequences. Residues conserved between all the analyzed proteins are shown (red background). The conserved DE residues (green asterisk, *) are indicated by a green box. The DUF4091 region is underlined.

Next, the β-NGA candidate sequences *dssm_2* and *dssm_3* were selected and heterologously expressed in *Escherichia coli*. The encoding sequences lacking the predicted signal peptides (Supplementary Data S1, bold and underlined) were cloned into an expression vector. Although an expression construct of *dssm_2* failed to yield a soluble protein, the enzyme encoded by *dssm_3* was solubilized, and a purified protein was successfully obtained (renamed Protein ID: NgaDssm). Assays using various *p*NP-substrates revealed that NgaDssm was active on GalNAc-β-*p*NP, but not on GlcNAc-β-*p*NP or GalNAc-α-*p*NP, indicating that the enzyme possessed exo-β-NGA but not exo-β-HEX activity. Furthermore, the enzyme hydrolyzed Galβ1-3GalNAc-β-*p*NP but not Gal-β-*p*NP, demonstrating an additional disaccharide-releasing endo-β-NGA activity (Table 1). These findings suggest that NgaDssm is a novel endo/exo-β-NGA.

**Table 1.**
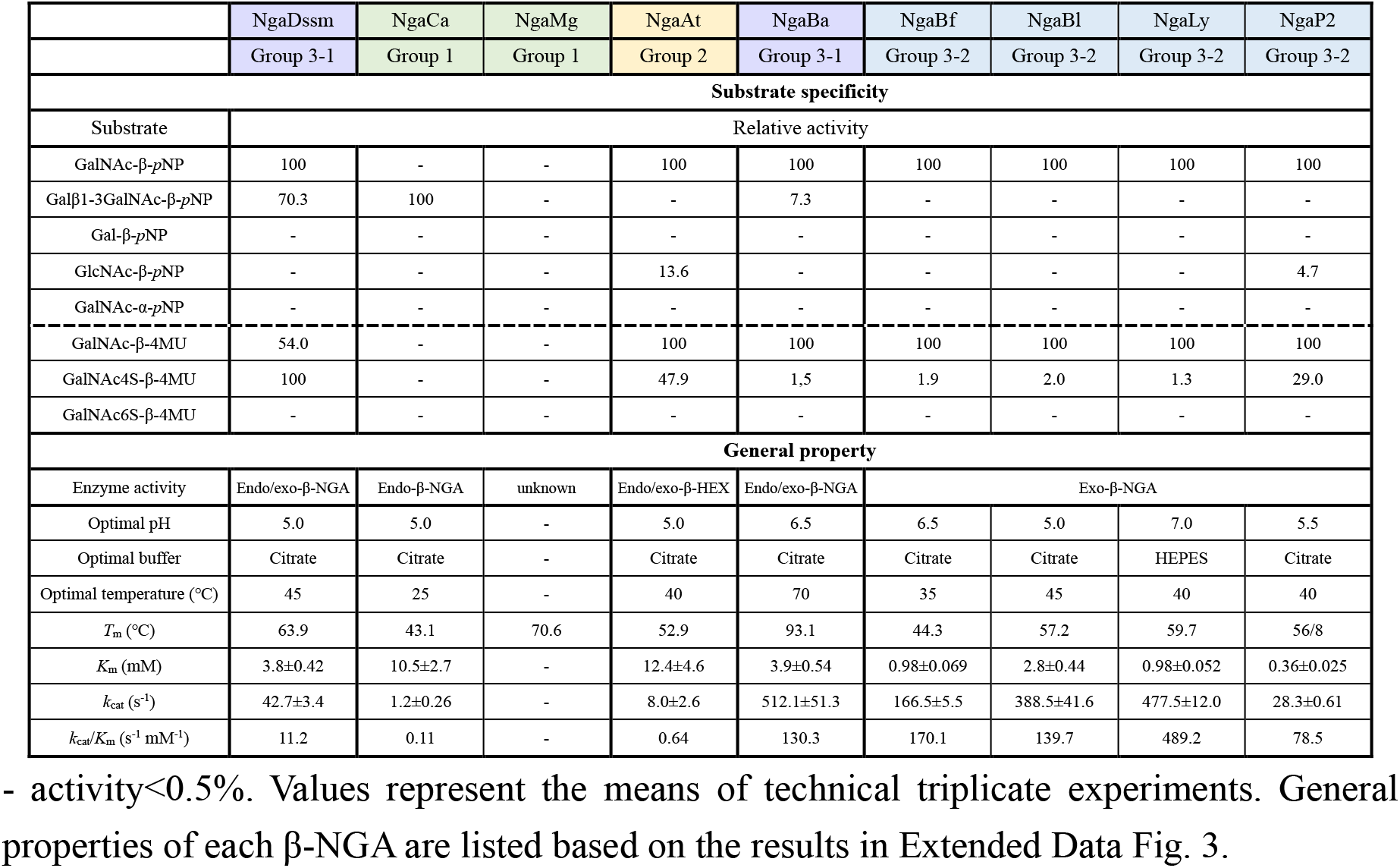
Substrate specificities and general properties of β-NGAs.

|  | NgaDssm | NgaCa | NgaMg | NgaAt | NgaBa | NgaBf | NgaBl | NgaLy | NgaP2 |
| --- | --- | --- | --- | --- | --- | --- | --- | --- | --- |
|  | Group 3-1 | Group 1 | Group 1 | Group 2 | Group 3-1 | Group 3-2 | Group 3-2 | Group 3-2 | Group 3-2 |
| Substrate specificity |  |  |  |  |  |  |  |  |  |
| Substrate | Relative activity |  |  |  |  |  |  |  |  |
| GalNAc-β-pNP | 100 | - | - | 100 | 100 | 100 | 100 | 100 | 100 |
| Galβ1-3GalNAc-β-pNP | 70.3 | 100 | - | - | 7.3 | - | - | - | - |
| Gal-β-pNP | - | - | - | - | - | - | - | - | - |
| GlcNAc-β-pNP | - | - | - | 13.6 | - | - | - | - | 4.7 |
| GalNAc-α-pNP | - | - | - | - | - | - | - | - | - |
| GalNAc-β-4MU | 54.0 | - | - | 100 | 100 | 100 | 100 | 100 | 100 |
| GalNAc4S-β-4MU | 100 | - | - | 47.9 | 1.5 | 1.9 | 2.0 | 1.3 | 29.0 |
| GalNAc6S-β-4MU | - | - | - | - | - | - | - | - | - |
| General property |  |  |  |  |  |  |  |  |  |
| Enzyme activity | Endo/exo-β-NGA | Endo-β-NGA | unknown | Endo/exo-β-HEX | Endo/exo-β-NGA | Exo-β-NGA | | | |
| Optimal pH | 5.0 | 5.0 | - | 5.0 | 6.5 | 6.5 | 5.0 | 7.0 | 5.5 |
| Optimal buffer | Citrate | Citrate | - | Citrate | Citrate | Citrate | Citrate | HEPES | Citrate |
| Optimal temperature (°C) | 45 | 25 | - | 40 | 70 | 35 | 45 | 40 | 40 |
| $T_m$ (°C) | 63.9 | 43.1 | 70.6 | 52.9 | 93.1 | 44.3 | 57.2 | 59.7 | 56/8 |
| $K_m$ (mM) | 3.8±0.42 | 10.5±2.7 | - | 12.4±4.6 | 3.9±0.54 | 0.98±0.069 | 2.8±0.44 | 0.98±0.052 | 0.36±0.025 |
| $k_{cat}$ (s<sup>-1</sup>) | 42.7±3.4 | 1.2±0.26 | - | 8.0±2.6 | 512.1±51.3 | 166.5±5.5 | 388.5±41.6 | 477.5±12.0 | 28.3±0.61 |
| $k_{cat}/K_m$ (s<sup>-1</sup> mM<sup>-1</sup>) | 11.2 | 0.11 | - | 0.64 | 130.3 | 170.1 | 139.7 | 489.2 | 78.5 |

### Phylogenetic diversity of β-NGAs

The uncharacterized domain DUF4091 was conserved in the GH123 exo-β-NGAs and a novel endo/exo-NGA gene sequence. Consequently, we utilized DUF4091 as a query to further identify β-NGA genes. We retrieved 734 sequences containing DUF4091 from the Pfam protein family database. The catalytic DE motif was highly conserved (94%), and thus, majority of these genes were expected to encode enzymes possessing β-NGA activity. A phylogenetic tree of these sequences, along with the three known GH123 exo-β-NGA genes and the two deep-sea β-NGA candidates (Supplementary Data S2), identified four major groups: GH123, Group 1, Group 2, and Group 3 (Fig. 1c). Group 3 was further segregated into two subgroups (Group 3-1 and Group 3-2) based on the phylogenetic analysis, with NgaDssm belonging to Group 3-1 (Fig. 1c, d).

To examine the β-NGA sequence alignment and activity distributed among all three domains of life, 14 representative sequences of each novel group were selected from model plants, archaea, and various bacterial phyla, as follows: for Group 1, *Cohnella abietic* from Bacillota (NgaCa), *Meiothermus granaticius* from Deinococcota (NgaMg), *Nostoc punctiforme* (NgaNp), *Clyndrospermum stagnale* (NgaCs), and *Stanieria cyanosphaera* (NgaSc) from Cyanobacteria; for Group 2, *Arabidopsis thaliana* (NgaAt), *Glycine max* (NgaGm), and *Oryza sativa* (NgaOs) from plants; for Group 3-1; *Candidatus Bathyarchaeia archaeon* B24 from Thermoproteota in Archaea from the hydrothermal vent microbiome (NgaBa) and *Chitinophaga pinensis* from Bacteroidota (NgaCp); for Group 3-2; *Brachybacterium faecium* (NgaBf) and *Bifidobacterium longum* subsp. *infantis* (NgaBl) from Actinomycetota, *Lacticaseibacillus yichunensis* (NgaLy), and *Paenibacillus* sp. TS12 (NgaP2) from Bacillota (Supplementary Data S2 and 3). Overall, the sequence identities between each group of 14 candidate genes and the three GH123 genes were low (12–26%) based on sequence alignment (Extended Data Fig. 2a), and only nine amino acids were entirely conserved (Fig. 1d, red background). DUF4091 was a comparatively well-conserved region, comprising four of the nine strictly conserved amino acids, with its sequence identity being at least 10% higher than that of the full-length sequence (Extended Data Fig. 2b, c).

### Substrate specificity of β-NGA candidates

Recombinant expression vectors for the new candidate genes (except for NgaGm and NgaOs) were constructed to assess the activity of the identified enzymes (Supplementary Data S3). Although four expression constructs (NgaNp, NgaCs, NgaSc, and NgaCp) failed to yield soluble proteins, the remaining eight (NgaCa, NgaMg, NgaAt, NgaBa, NgaBf, NgaBl, NgaLy, and NgaP2) successfully expressed soluble enzymes (Extended Data Fig. 2a, red letter) and were subjected to protein purification for subsequent enzymatic assays (Extended Data Fig. 3a, Table 1).

The substrate specificity of the expressed enzymes was assayed using various synthetic substrates (Table 1). Surprisingly, Group 1 NgaCa displayed strictly endo-type enzymatic activity and acted solely on Galβ1-3GalNAc-β-*p*NP. The other Group 1 protein, NgaMg, demonstrated no activity against any of the tested substrates.

In Group 2, NgaAt was active against GalNAc-β-*p*NP and GlcNAc-β-*p*NP, but not against Galβ1-3GalNAc-β-*p*NP, indicating that it possessed exo-β-HEX activity. However, NgaAt did not share notable sequence similarity with any of the three types of exo-β-HEXs (HEXO1–3) classified into the GH20 family from *A. thaliana* (Extended Data Fig. 4a, b) (26, 27). Thus, NgaAt is a novel family of exo-β-HEX.

NgaBa in Group 3-1 was active on both GalNAc-β-*p*NP and Galβ1-3GalNAc-β-*p*NP, similar to NgaDssm, suggesting that members of Group 3-1 share endo/exo-type β-NGA activity. In Group 3-2, NgaBf, NgaBl, and NgaLy were active only on GalNAc-β-*p*NP, highlighting their strict exo-β-NGA functionality. NgaP2 was active against GalNAc-β-*p*NP and had weak activity against GlcNAc-β-*p*NP. Collectively, these results showed that the enzymes in each group exhibit a characteristic endo- and/or exo-type cleavage mode and substrate specificity.

NgaP from GH123 is an excellent tool for detecting sulfatase deficiency, as it does not act on substrates sulfated at positions 4 or 6 of GalNAc (28), and has been successfully used for screening of mucopolysaccharidosis (a metabolic disorder caused by the accumulation of mucopolysaccharides) in newborns (28, 29). We evaluated whether the enzymes identified herein possessed GalNAc4S or GalNAc6S cleavage activity. Using GalNAc4S-β-4MU and GalNAc6S-β-4MU (Table 1), we observed that NgaDssm, NgaAt, and NgaP2 acted on GalNAc4S-β-4MU. In particular, NgaDssm displayed approximately two-fold higher activity against GalNAc4S-β-4MU than against GalNAc-β-4MU.

### Generic properties of the novel β-NGAs

Next, we evaluated the enzyme characteristics (Table 1, Extended Data Fig. 3). The optimal pH for the eight novel enzymes was in the range 5.0–7.0. NgaBa depicted a particularly broad optimal pH range, with relative activity above 90% at pH 5.5–7.5. Metal ions generally did not affect enzyme activity, except that of NgaLy, which was inhibited by Ni^2+^, Co^2+^, and Zn^2+^ (Table 1, Extended Data Fig. 3b, c). Overall, the enzymes exhibited moderate temperature optima (25–45℃) except for NgaBa, an enzyme from a hydrothermal vent microbiome (30), which displayed an extremely high optimal temperature (70℃) and *T*_m_ value (93.1℃) (Extended Data Fig. 3d, e). Thus, NgaBa is the first thermostable β-NGA reported to date. The *K*_m_ and *k*_cat_ of each enzyme for the most preferred substrate (GalNAc-β-*p*NP or Galβ1-3GalNAc-β-*p*NP) was also examined (Extended Data Fig. 3f), wherein NgaLy possessed the highest *k*_cat_/*K*_m_ value.

### Substrate specificity for oligosaccharides

The substrate specificity of each enzyme was explored using various oligosaccharides as substrates (Extended Data Fig. 5a). NgaCa and NgaMg from Group 1 and NgaDssm from Group 3-1 surprisingly did not act on GA1 and Gb5 oligosaccharides, although these two oligosaccharides shared the non-reducing end structure Galβ1-3GalNAc-β- with Galβ1-3GalNAc-β-*p*NP (Extended Data Fig. 5b, d). NgaAt acted on GalNAcβ1-3Gal but not on GalNAcβ1-4Gal (Extended Data Fig. 5c). By contrast, NgaBa was functional against both GA1 and Gb5 (Extended Data Fig. 5d, right lanes 2 and 5), displaying a more rapid digestion of GA1 over that of GA2 (Extended Data Fig. 5d, right lanes 2 and 4). The endo-activity of NgaBa against β-GalNAc located inside the oligosaccharide was stronger than its exo-activity against β-GalNAc at the non-reducing ends, but this preference was reversed when *p*NP-substrates were used. Moreover, NgaBa more effectively degraded Gb5 than Gb4 (endo-> exo-activity) and displayed enzymatic activity against Galβ1-3GalNAc-β- and GalNAcα1-3GalNAc-β-(Extended Data Fig. 5d). The NgaBf, NgaBl, NgaLy, and NgaP2 activities were similar to those of GH123 exo-β-NGAs, as they broke down only linear oligosaccharides with β-GalNAc at the non-reducing terminus (Extended Data Fig. 5e). The results for NgaBf and NgaLy indicated that these enzymes preferred GalNAcβ1-4Gal to GalNAcβ1-3Gal. As only a few β-GalNAc-containing oligosaccharides are commercially available, the natural substrates of NgaCa, NgaMg, and NgaDssm were not identified.

### X-ray crystal structures of the β-NGA from novel groups

Although the novel β-NGAs and GH123 exo-β-NGAs were divided into four major groups based on the phylogenetic analysis, their predicted overall structure was similar. Based on substrate preferences, several of these enzymes considerably differed from GH123 exo-β-NGAs in terms of substrate specificity. To understand the relationship between structural variation and substrate specificity, crystallization screening was performed on all novel β-NGAs, and X-ray crystallographic analyses of apo- and/or ligand-bound forms were successfully performed on the following enzymes (NgaCa [Group 1], NgaAt [Group 2], NgaDssm [Group 3-1], NgaLy, and NgaP2 [Group 3-2]; nine forms of five enzymes in total) (Fig. 2 and Supplementary Tables 1–5). The overall structure of these enzymes is similar to that of CpNga123 and BvGH123, which consist of a β-sandwich domain at the N-terminus and a (β/α)_8_-barrel domain encompassing the catalytic region (Fig. 2a, Supplementary Figs. 1–5). The structure of NgaDssm (Group 3-1) bore striking resemblance to that of NgaLy and NgaP2 (Group 3-2) (root-mean-square distance of Cα atoms [rmsd] = 1.5–2.3 Å), while the structures of NgaLy and NgaP2 are identical (rmsd = 1.0 Å) (Extended Data Fig. 6a). The additional N-terminal domain consisting of approximately 80 amino acids, found only in Group 2 enzymes, was not modeled in the crystal structure of NgaAt due to disorder (Supplementary Fig. 2).

**Fig. 2.**
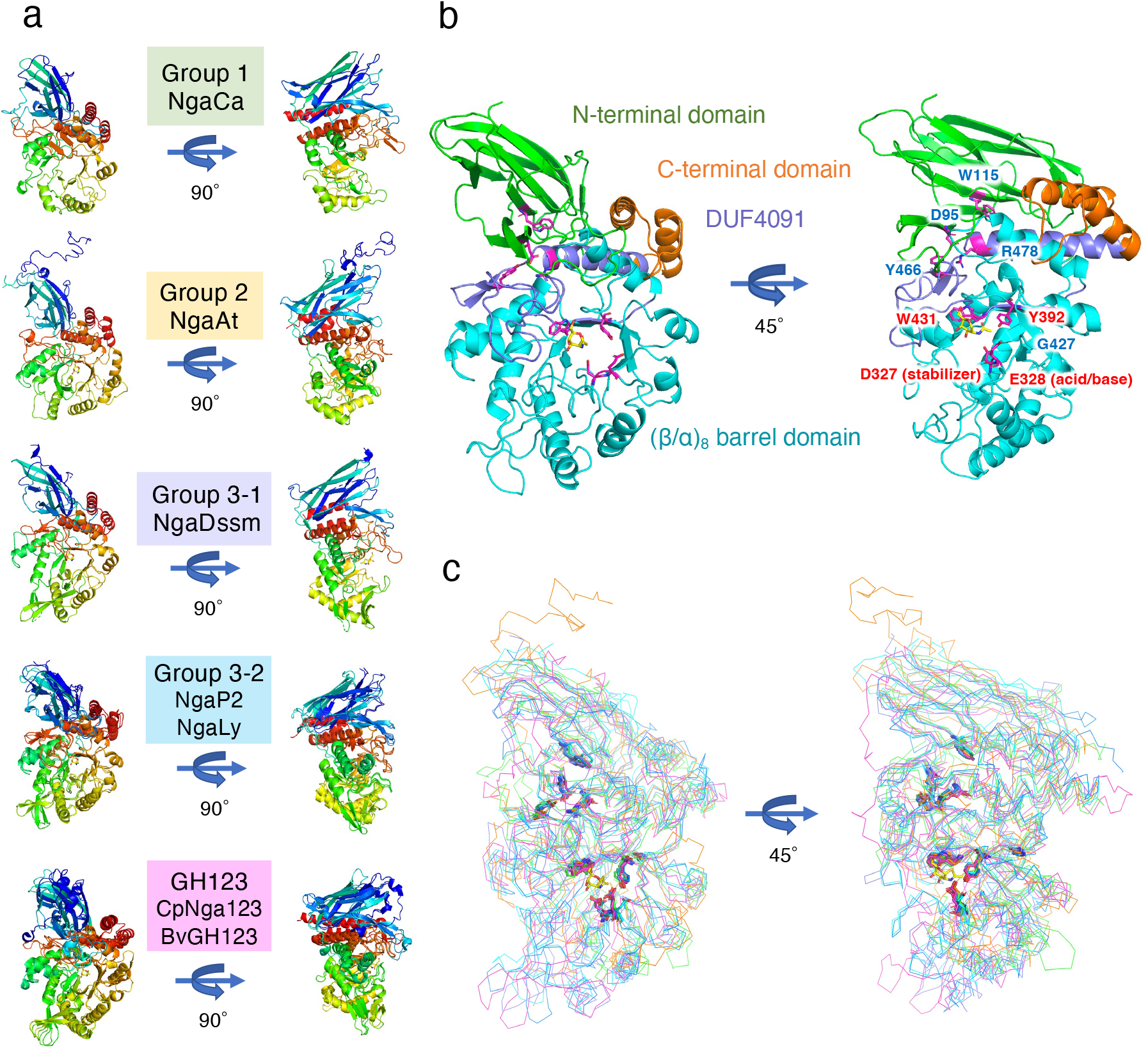
Overall structures and superposition of β-NGAs. **a**, Structures of β-NGAs. The DE motif is shown in stick format. **b**, The structures of NgaLy. β-Sandwich domain at the N-terminus, (β/α)_8_-barrel domain, DUF4091 and C-terminal domain are shown in green, cyan, purple, and orange, respectively. The conserved amino acids are shown as magenta sticks. Ligands are placed at the substrate-binding site and shown as yellow sticks. **c**, Superposition of the Group 1, Group 2, Group 3 and GH123 structures. Conserved amino acids are plotted on the structures as sticks.

Among Groups 1, 2, and 3 and GH123, the DE motif is located at the same position, suggesting that these enzymes adopt the substrate-assisted catalysis reported for GH123 exo-β-NGAs (Fig. 2b). The conserved DUF4091 domain in these enzyme groups is located at the innermost position of the enzyme between the N-terminal domain and the (β/α)_8_-barrel domain (Fig. 2b, purple). This suggested that DUF4091 most likely plays a role in defining the location of N-terminal and (β/α)_8_-barrel domains as a “central pillar”. Intriguingly, nine amino acid residues conserved among the groups (Fig. 1d, red background) are found at the same positions in the structure (Fig. 2c). Among the conserved amino acids, D95, W115, Y466, and R478 (amino acid numbers of NgaLy [Extended Data Fig. 6b]) are located at the interface between the N-terminal domain and DUF4091 (Fig. 2b, right). D95 and W115, located at the N-terminal domain, fit into the surface pocket of DUF4091, and the side chain of D95 formed hydrogen bonds with Y466 and R478 located at DUF4091 (Extended Data Fig. 6c). Therefore, D95, W115, Y466, and R478 are potentially important for maintaining structural integrity as they link the N-terminal and (β/α)_8_-barrel domains (Fig. 2b, blue letter). G427 is positioned at the entry point for the eighth β-sheet into the (β/α)_8_ barrel domain, with the space barely the size of the glycine residue (Extended Data Fig. 6d). The DE motif (D327 and E328), Y392, and W431 are located around the substrate and are involved in substrate recognition (Fig. 2b, red letter). Previous studies have reported that point mutations in the DE motif decrease catalytic activity (9, 12). Therefore, we examined the remaining seven conserved amino acids and constructed corresponding alanine mutants (Extended Data Fig. 6e). The point mutants D95A, W115A, Y466A, and R478A were expressed in *E. coli*, but all proteins precipitated. For G427, a Val mutant was also constructed based on structural information that suggested the conversion of Gly to Ala would be tolerated (Extended Data Fig. 6d). In this case, a very small amount of solubilized enzyme was obtained for G427A but not for G427V (Extended Data Fig. 6e, blue letter). The Y392A and W431A mutants were purified as soluble enzymes (Extended Data Fig. 6e, red letter), but their catalytic activity was considerably reduced (Extended Data Fig. 6f). These findings imply that amino acid substitutions severely impact the structural stability (D95, W115, Y466, R478, and G427) and substrate recognition (Y392 and W431) of the enzyme.

### Essential structural elements for endo- and exo-specificity

We compared the overall structure and substrate-binding sites among the groups to identify the structural elements governing endo- and exo-specificity of these enzymes (Fig. 3). NgaCa in Group 1 had the simplest structure among the enzymes analyzed (Fig. 3a, Supplementary Fig. 1). The active site of NgaCa possesses a cleft shape that allows oligosaccharides to pass through to the −2 subsite, enabling oligosaccharide binding and endo-type β-NGA activity. This structural feature indicated that Group 1 enzymes act on the inner regions of longer sugar chains in addition to disaccharides from the non-reducing end (e.g., Galβ1-3GalNAc-β-*p*NP).

**Fig. 3.**
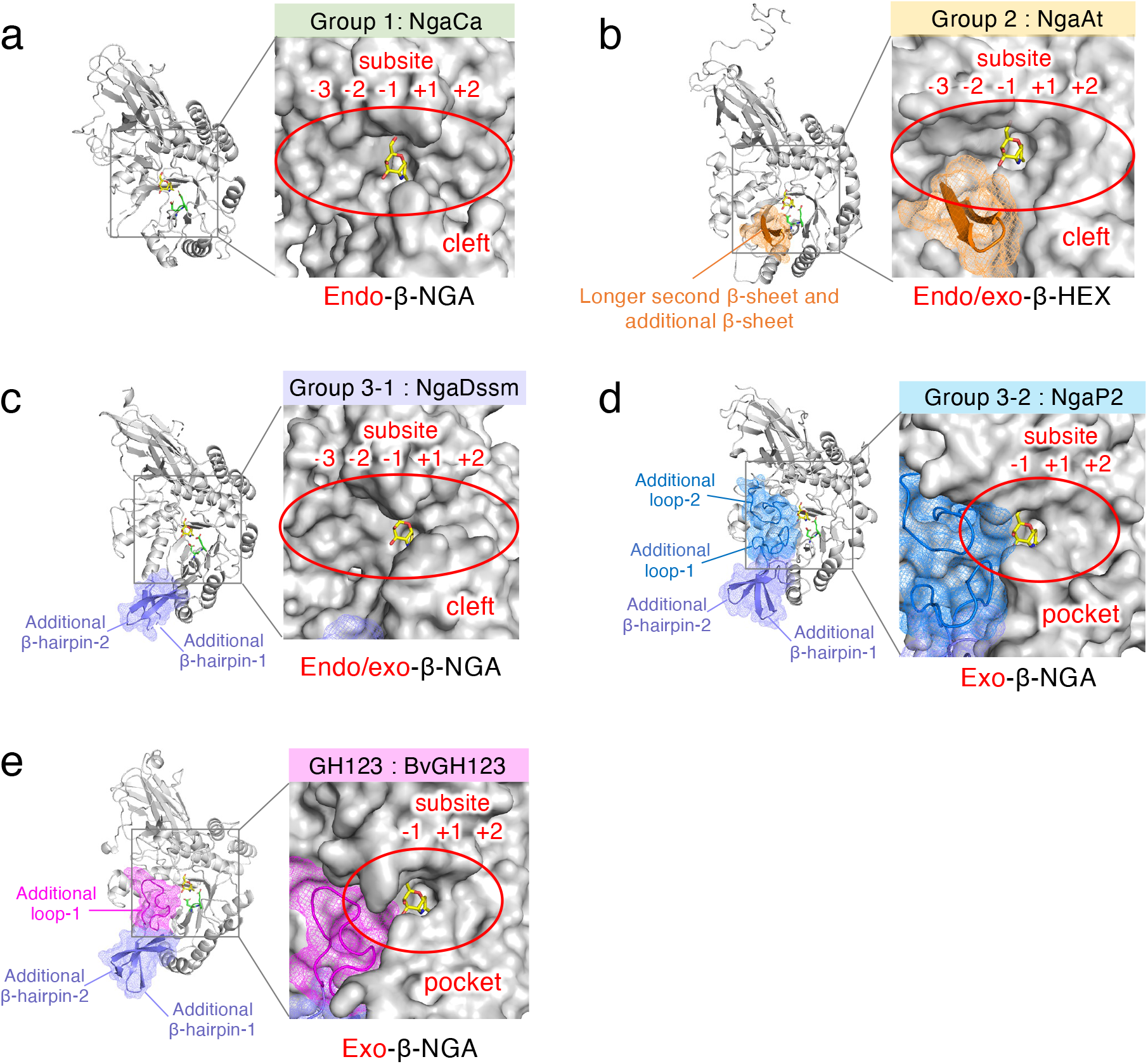
Substrate-binding area of β-NGAs. The overall structure of β-NGAs is shown as a cartoon (left) and the molecular surface of the substrate-binding site of β-NGA complexed with the ligand (right). The core structure (Group 1) is shown in gray (**a**), while additional regions characteristic of the other groups are shown in (**b**) orange (Group 2), (**c**) purple (Group 3-1), (**d**) blue (Group 3-2), and (**e**) magenta (GH123). The DE motif and ligand are indicated by green and yellow sticks, respectively.

The active site of NgaAt in Group 2 also possesses a cleft shape (Fig. 3b), and the second β-sheet of the (β/α)_8_ barrel is longer than that found in the other enzymes, followed by an additional β-sheet (Fig. 3b and Supplementary Fig. 2, orange). Consequently, the space around the 3-OH group of β-GalNAc in the −2 subsite is narrower than that in Group 1. Therefore, NgaAt was presumed to act on Galβ1-4GalNAcβ- but not on Galβ1-3GalNAcβ-. Since Galβ1-4GalNAc-β-*p*NP was commercially unavailable, Galβ1-4GlcNAc-β-*p*NP was used as an alternative substrate to test this hypothesis, and we observed that Galβ1-4GlcNAc-β-*p*NP, but not Galβ1-3GalNAc-β-*p*NP, was degraded (Extended Data Fig. 4c). Thus, NgaAt demonstrates more diverse activity than expected and possesses endo/exo-β-HEX functionality, making it the first β-HEX with this characteristic.

The active site of the endo/exo-type NgaDssm from Group 3-1 has a cleft structure that enables passage to the −2 subsites, as in NgaCa, but it also possesses two additional β-hairpins from the second and third β-sheets of the (β/α)_8_ barrel domain (Fig. 3c and Supplementary Fig. 3, purple). Moreover, NgaP2 (Group 3-2) has two loops above and below the cleft from the second and eighth β-sheets of the (β/α)_8_ barrel domain (Fig. 3d and Supplementary Fig. 4, blue). The strict exo-type activity of Group 3-2 can be explained by these two loops that completely block the −2 subsite, forming a pocket-like architecture at the substrate-binding site and thus preventing substrate entry.

Similarly, BvGH123 in GH123 also has two additional β-hairpins and one extended loop from the second β-sheet of the (β/α)_8_ barrel domain, which blocks the −2 subsite side of the cleft, yielding a pocket-like conformation (Fig. 3e and Supplementary Fig. 5, magenta).

These findings revealed that substrate-binding sites with cleft structures exhibit endo-type activity, while those with pocket-like architectures show exo-type activity.

### Substrate recognition and catalytic mechanism of β-GalNAc-acting enzymes

We further examined detailed substrate recognition mechanisms using structural analysis of complexes with GalNAc-thiazoline, an analog of the oxazolinium intermediate and a potent inhibitor of enzymes utilizing substrate-assisted catalysis (31, 32) (Fig. 4, Supplementary Fig. 6). With the exception of NgaCa, the structures of the NgaAt, NgaDssm, and NgaP2 complexes with GalNAc-thiazoline were successfully characterized (Supplementary Tables 2–4), and a subsequent docking model was engineered for NgaCa.

**Fig. 4.**
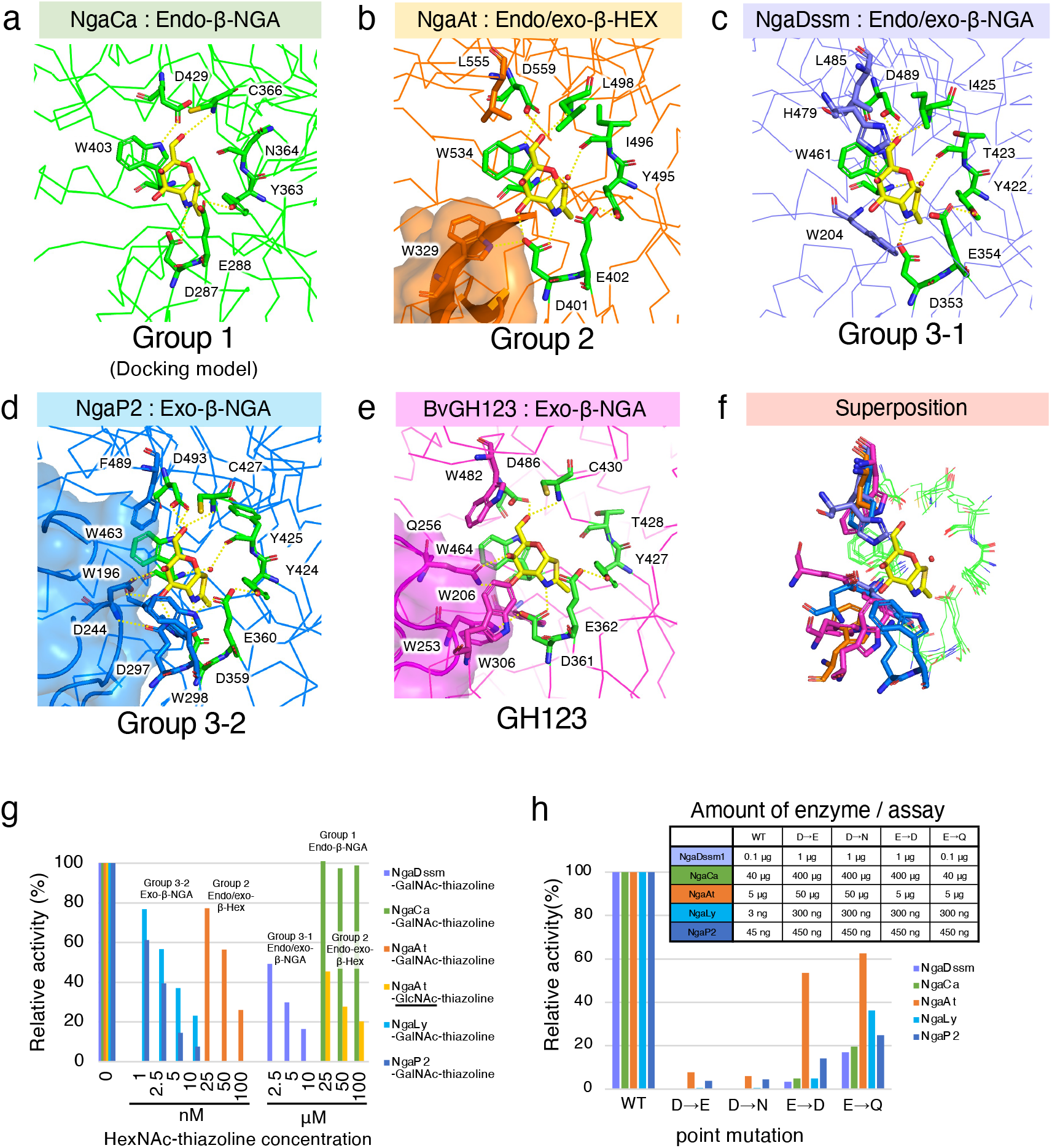
Substrate recognition and catalytic mechanism. **a–e**, Active-site structure of each enzyme. **a**, Docking model of NgaCa with GalNAc-thiazoline. **b–e**, Crystal structure of each enzyme complexed with GalNAc-thiazoline. Hydrogen bonds are indicated by dotted lines. **f**, Superimposition of active sites in Groups 1, Group 2, Group 3 and GH123. **g**, Inhibition of β-NGA activity by GalNAc-thiazoline and GlcNAc-thiazoline. **h**, Point mutation analysis of the DE motif. The amounts of enzyme used in the assay are shown in the top-right table.

The docking model of NgaCa (Group 1) demonstrated that seven amino acid residues recognize GalNAc-thiazoline (Fig. 4a, green residues). In NgaAt (Group 2), apart from these seven residues, W329 establishes a hydrogen bond with GalNAc-thiazoline, and L555 contributes to substrate positioning (Fig. 4b, orange residues). Similarly, in NgaDssm (Group 3-1), the H479 residue forms a hydrogen bond, while residues W204 and L485 are involved in substrate positioning (Fig. 4c, purple residues). These results imply that Group 2 and Group 3-1 members possess a higher affinity for β-GalNAc than those of Group 1. Furthermore, complex binding modes were observed for Group 3-2 and GH123. For NgaP2 (Group 3-2), in addition to the seven residues discussed for NgaCa, three (D244, D297, and W298) and two (W196 and F489) residues are involved in hydrogen bond formation and substrate positioning, respectively (Fig. 4d, blue residues). Similarly, in BvGH123 (GH123), two (W253 and Q256) and three (W206, W306, and W482) residues are involved in these processes (Fig. 4e, magenta residues). Compared to the amino acid positions of the endo-type enzyme (green sticks), the exo-type enzymes have additional substrate-binding amino acid residues around β-GalNAc, indicating stronger recognition from the −2 subsite side. The increased number of residues could potentially reinforce substrate recognition in Group 3-2 and GH123 enzymes (Fig. 4f). Additional structural analyses (comparison of crystal structure with AlphaFold2 predicted structure of NgaCa, comparison of apo1-form and apo2-form of NgaCa, and comparison of GalNAc-thiazoline-bond form and GlcNAc-thiazoline-/apo-form of NgaAt, NgaDssm, and NgaP2) data are illustrated in Extended Data Fig. 7.

To complete the enzymatic analysis, we examined GalNAc- or GlcNAc-thiazoline as inhibitors of these enzymes. In the assays using GalNAc-thiazoline, Group 3-2 exo-β-NGAs showed inhibitory activity at >1 nM concentrations (Fig. 4g, blue and light blue). By contrast, no inhibition was observed in Group 1 endo-β-NGA even at a concentration of 100 μM (Fig. 4g, green). Interestingly, the activity of Group 3-1 endo/exo-β-NGA was inhibited at concentrations 1000-fold higher than for exo-β-NGA (Fig. 4g, purple). NgaAt was inhibited by 25 nM GalNAc-thiazoline and by 25 μM GlcNAc-thiazoline (a 1000-fold higher concentration) (Fig. 4g, orange and yellow). These results corroborate the disparity in recognition abilities of GalNAc-thiazoline identified from the aforementioned structures.

### Analysis of catalytic residue mutants

Based on structural analysis, Asp and Glu of the DE motifs were recognized as the stabilizer of the 2-acetamido group and acid-base catalytic residue, respectively. The GalNAc-thiazoline-bound structures and inhibition assays indicated that these enzymes perform substrate-assisted catalysis. Therefore, a point mutation analysis of the DE motifs was conducted (Fig. 4h). Mutants with Asp-to-Glu/Asn alterations exhibited little to no activity even at 10-fold to 100-fold higher enzyme concentrations. Glu-to-Asp/Gln mutants demonstrated reduced enzymatic activities, and the trends were similar to those observed for other enzymes performing substrate-assisted catalysis (9, 12).

### Elucidation of the catalytic mechanism using NMR

GH123 enzymes are exo-β-NGAs, while enzymes from Groups 1 and 3-1 are endo- and endo/exo-type β-NGAs, respectively. To further confirm the substrate-assisted catalysis mechanism of these enzymes, we investigated the reaction products by NMR using Galβ-1-3GalNAcβ-*p*NP as a substrate (Extended Data Figs. 8 and 9, Supplementary Figs. 7 and 8, Supplementary Tables 6 and 7) and monitored the stereochemistry of glycosidic bond hydrolysis (between the GalNAc and *p*NP moieties) by ^1^H NMR. The anomeric hydrogen signal of Galβ-1-3GalNAcβ-*p*NP (between the GalNAc and *p*NP moieties) disappeared within 1 min for NgaDssm (Extended Data Fig. 8a) and 10 min for NgaCa (Extended Data Fig. 8b) and Galβ-1-3GalNAcβ appeared, while Galβ-1-3GalNAcα anomeric signals appeared after 10 and 30 min, respectively, due to mutarotation. These results indicate that NgaDssm and NgaCa are anomer-retaining enzymes, similar to GH123 enzymes.

## Discussion

We used deep-sea metagenomic sequences to discover a novel β-NGA, NgaDssm, which is the first β-NGA to possess dual endo/exo-type-β-NGA activity and low sequence similarity to known exo-β-NGAs. Prior studies have characterized three GH123 exo-β-NGAs (NgaP, CpNga123, and BvGH123) from land soil (9) and human gut bacteria (12, 13); however, no genes from other ecological niches have been reported. The deep-sea environment (below 200 m depth) is characterized by total darkness, low temperatures, and high pressure, with occasional high temperatures owing to geological formations, such as hydrothermal vents. The deep-sea environments are completely distinct from the terrestrial environment and remain unexplored owing to limited accessibility to samples. Therefore, the deep-sea microbiome is an attractive potential bioresource for screening undiscovered enzymes (23).

We further discovered novel endo-, endo/exo-, and exo-type β-NGAs, as well as endo/exo-β-HEXs, acting on β-GalNAc by analyzing deep-sea metagenomic sequences and public protein databases, and our comparative biochemical and structural analyses provide insights into the molecular evolution of β-GalNAc-targeting enzymes (Fig. 5). Despite their phylogenetic distance and different substrate specificities, these novel enzymes and GH123 β-NGAs are likely homologous proteins based on the presence of conserved residues in their sequences (Fig. 1d), which are also positionally retained in their structures (Fig. 2c), and the observation that mutations in conserved residues were not tolerated and resulted in destabilization of protein structure and elimination of substrate recognition (Extended Data Figs. 6e, f). Structural comparisons between members in Groups 1–3 and GH123 enzymes further supported that the four families have diversified their substrate specificity (endo-β-NGA for Group 1, endo/exo-β-HEX for Group 2, endo/exo-β-NGA, exo-β-NGA for Group 3, and exo-β-NGA for GH123) through the accumulation of point mutations and insertional sequences (Figs. 3 and 4). These data suggested a monophyletic evolutionary history of β-NGAs from the prototype enzymes in Group 1 β-NGAs (Fig. 5).

**Fig. 5.**
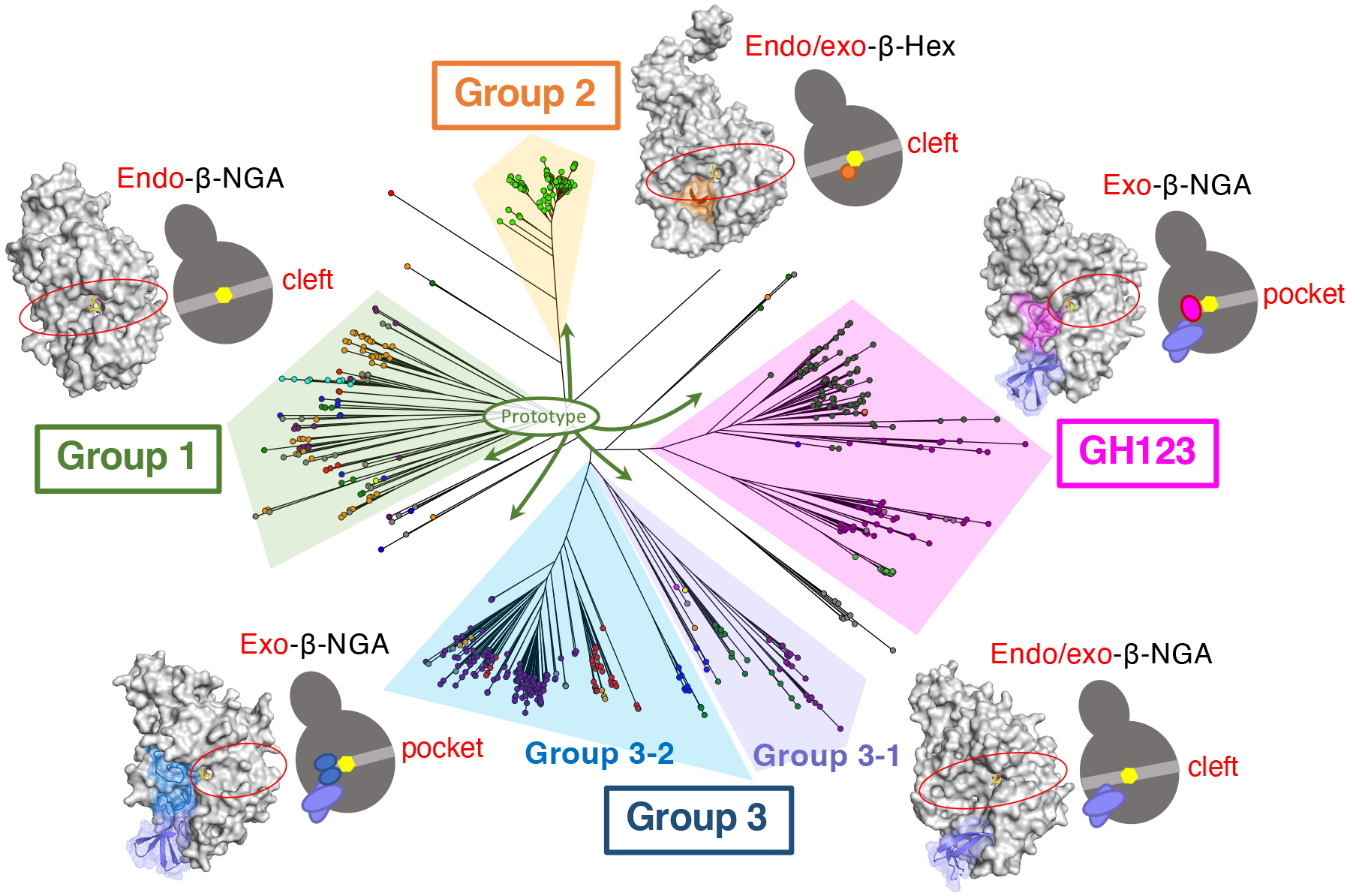
Phylogenetic tree and structural features of β-NGAs. The overall structure of β-NGAs is illustrated as a surface model with an accompanying schematic representation. The ligand is indicated by a yellow stick. The Group 1 structure is depicted in gray as the basic structure. Additional regions characteristic of other groups are highlighted in orange (Group 2), purple (Group 3-1), blue (Group 3-2), and magenta (GH123).

In the CAZy database, several GH families are grouped into “clans” according to the conservation of catalytic residues, common reaction mechanisms, and structural similarities (11, 33). Consistent with these criteria, we suggest a new clan composed of the enzyme groups discovered in this study (Groups 1, 2, and 3) and GH123. Among the new enzymes identified herein, only NgaAt (Group 2) is a β-HEX that acts on β-GalNAc and β-GlcNAc. We speculate that a prototype enzyme of the new putative GH clan, including GH123 was a β-NGA specific for β-GalNAc, and the β-HEX enzymes of Group 2 are proposed to be divergent from the structurally simpler β-NGA. By contrast, GH20, mainly comprising β-HEX enzymes, belongs to clan GH-K, together with GH18 and GH85. All GH-K enzymes act on β-GlcNAc bonds (e.g., exo- and endo-β-*N*-acetylglucosaminidase, chitinase, lacto-*N*-biosidase, and β-hyaluronidase). Thus, GH-K enzymes seem to originate from a structurally simple β-*N*-acetylglucosaminidase, and GH20 β-HEX appears to belong to a family that diverged from the prototypical β-*N*-acetylglucosaminidase. Therefore, since β-HEX acts on β-GalNAc and β-GlcNAc, we suggest that β-HEX has two enzyme lineages with different evolutionary pathways.

In a previous study on β-NGAs from GH123, the function of DUF4091 has never been reported (12, 13). However, our domain-based exploration of β-NGA genes, based on GH123 members in the deep-sea metagenomic data, led to the identification of novel β-NGA types containing DUF4091. Mutation experiments also highlighted that DUF4091 is indispensable for the stability of β-NGAs. The functional annotation of CDSs is typically conducted based on the similarity of the entire sequence to known protein sequences. A domain-based search (typically based on hidden Markov models) is an alternative method for identifying homologous sequence regions, even if the overall similarity is low. Examining CAZy families with consideration of co-occurring functionally unknown domains will likely lead to the subsequent discovery of additional novel enzymes.

To investigate the possible biological functions of the novel β-NGAs, we examined neighboring genes and protein-protein interactions using the STRING database (34) (Extended Data Fig. 10, Supplementary Data S4 and S5). The enzymes identified herein are potentially involved in the regulation of lipoprotein (NgaCa), the capsular membrane (NgaCp), and *O*-antigen (NgaCs), as well as in the degradation of glycans in the periplasm (NgaBf). Therefore, β-NGAs may exhibit more diverse functions than solely the degradation and utilization of glycans, which GH123 β-NGA is thought to be responsible for and could have roles in various GalNAc-mediated biological processes.

Since the majority of prokaryotes in nature remain uncultured, it is crucial to explore the functional and genetic diversity of glycosidases in uncultured microorganisms through metagenomic analysis in order to gain a comprehensive understanding of glycan-mediated phenomena. The novel glycosidases hold potential for various industrial applications, including the utilization of these enzymes as biocatalysts for the production of functional oligosaccharides, glycan structure analysis, and disease diagnosis. Therefore, the discovery of novel glycosidases is crucial for both the basic and applied sciences. The comprehensive exploration and characterization of the diverse β-NGAs presented significantly enhance our understanding of their biological functions and their evolutionary history. These findings present a new approach to enunciating the evolutionary history of not only β-NGA but also many glycan-related enzymes. Further elucidation of the correlation between structures and diverse functions determined during evolution would also have significant implications for the structural basis of enzyme design for the engineering of new enzymes.

## Methods

### Sediment sampling and metagenomic sequencing

Abyssal sediment core was collected using a gravity corer on a remotely operated vehicle *ABISMO* (35), at station IOB (located 29.2746° N, 143.7673° E, at a depth of 5747 m below sea level) during a cruise of the ship R/V *Kairei* KR11-11 (December 2011) owned by the Japan Agency for Marine-Earth Science and Technology (JAMSTEC). The acquired sediment core was immediately subsampled onboard and stored at −80°C for molecular biology analyses (36).

Approximately 5 mL of frozen subsampled sediments from four sections (extracted from the depth ranges 0–8, 13–23, 53–63, and 113–123 cm below the seafloor) were used for metagenomic analysis. The sections were selected based on the geochemical profile and microbial community composition of the sediment core, as previously reported (36). Environmental DNA was extracted using the DNeasy PowerMax Soil Kit (QIAGEN, Hilden, Germany), per manufacturer protocol, with the following minor modification to increase DNA yield: cells were agitated twice for 10 min each after incubation at 65°C. Sequence libraries were prepared using the KAPA HyperPrep Kit (KAPA Biosystems) or the Ovation SP+ Ultralow Library System (NuGEN Technologies, San Carlos, CA, USA), as previously described (37). Library pools were mixed with Illumina PhiX control libraries and sequenced using the Illumina MiSeq or HiSeq platforms (Illumina, San Diego, CA, USA) at JAMSTEC or Macrogen (Seoul, South Korea).

### Bioinformatics

For raw metagenomic sequence data, both ends of the reads containing low-quality bases (Phread quality score <20) and adapter sequences were trimmed using TrimGalore (https://github.com/FelixKrueger/TrimGalore) with default settings. Sequencing reads derived from the PhiX genome were removed using Bowtie2 (38). Low complexity sequences or those shorter than 100 bp were discarded using PRINSEQ++ (39). The remaining high-quality paired-end reads of each sample were individually assembled *de novo* using metaSPAdes (40). Full-length coding sequences (CDSs) in the contigs were predicted using Prodigal (41) in the anonymous mode (‘p meta’ setting). Protein domain annotations of the CDSs were achieved through HMMER (42) against Pfam (version 35.0) (43) and dbCAN2 (version v10) (44) with a cutoff domain e-value of ≤ 1E-3. All CDSs assigned to GH123 and measuring > 400 bp in length were retrieved as β-NGA candidates and used for further analysis.

In addition to metagenomic CDSs, those with > 400 bp and architecture similar to known β-NGA genes (containing the DUF4091 [PF13320] domain in the C-terminal region) were retrieved from Pfam (version 35.0). A phylogenetic tree was constructed using MAFFT 1. (45) with default settings and FastTree2 (46) with JC+CAT models. ClustalOmega (47) and the ColabFold software (48) were employed for sequence alignment and protein structure prediction, respectively.

### Construction of β-NGA expression vectors

The selected β-NGA candidates were artificially synthesized by codon optimization for recombinant expression in *E. coli* using Strings DNA Fragments Synthesis Service (Thermo Fisher Scientific, Waltham, MA, USA) (Supplementary Data S3). The signal peptides in the amino acid sequences (Supplementary Data S1 and S3, highlighted with bold and underlined font) were predicted using SignalP 5.0 (49) and were removed from the N-terminal side of NgaDssm, NgaNp, NgaCs, NgaSc, and NgaCp. The gene encoding NgaP2 was directly cloned from the genomic DNA of *Paenibacillus* sp. TS12 via PCR amplification. These synthetic genes were designed with additional sequences at both terminals to facilitate amplification using a common primer set (Supplementary Data S6). The genes were cloned into the pET-47b(+) expression vector (Merck KGaA, Darmstadt, Germany) using the In-Fusion HD Cloning Kit (Takara Bio, Shiga, Japan). Mutagenesis was performed using the In-Fusion HD Cloning Kit. Primer sequences are listed in Supplementary Data S6.

### Expression and purification of recombinant β-NGA

The expression vector and recombinant mutant plasmids were used to transform *E. coli* BL21 Star (DE3) cells. The cells were cultured in 50 mL of medium A (LB medium containing 50 μg/mL of kanamycin), incubated at 37°C for 16 h with shaking, inoculated into 1–2 L of medium A, and incubated at 37°C for another 2–3 h with shaking. Protein expression was induced by addition of isopropyl β-D-1-thiogalactopyranoside (IPTG) to the culture at a final concentration of 0.1 mM. After additional culturing at 16°C for 16 h, cells were harvested by centrifugation (10,000 *g* for 10 min) and suspended in 50 mL of buffer A (20 mM HEPES-Na [pH 7.5], 150 mM NaCl, 5% [v/v] glycerol, 1 mM DTT, and 50 mM imidazole). Following sonication, cell debris was removed by centrifugation (13,000 *g,* 4 °C for 20 min) and passed through a 0.45-µm pore-sized GD/X syringe filter (Cytiva, Marlborough, MA, USA). The supernatant was subjected to chromatography using the AKTA Prime Chromatography System (Cytiva). The sample was loaded onto a 5 mL HisTrap HP column (Cytiva) at a flow rate of 2 mL/min. The column was then washed with buffer A. The His-tagged protein was eluted with Buffer B (20 mM HEPES-Na [pH 7.5], 150 mM NaCl, 5% [v/v] glycerol, 1 mM DTT, and 300 mM imidazole). The eluted fractions were pooled and dialyzed against Buffer C (20 mM HEPES-Na [pH 7.5], 150 mM NaCl, and 1 mM DTT). To cleave the His-Tag, the HRV3C protease was dialyzed at 4 °C for 16 h. The enzyme was further purified by ion-exchange [5-mL HiTrapQ column (Cytiva)] and size exclusion chromatography (SEC) [HiLoad 16/600 Superdex 200 pg column (Cytiva)]. The presence of the desired protein was confirmed by SDS-PAGE. The molecular weights of all β-NGAs estimated by SEC were consistent with the calculated molecular weights of their monomers.

### Enzyme assays

The activity of β-NGA candidates was determined using Assays I–III. In Assay I (*p*NP-β-GalNAc as a substrate), the reaction mixture comprised 50 nmol of *p*NP-β-GalNAc and an appropriate amount of the enzyme in 100 μL of a 100 mM optimal buffer solution. Following a 30-min incubation, the reaction was arrested by adding 100 μL of 1 M sodium carbonate, and the corresponding absorbance was measured at 405 nm. One unit of the enzyme was defined as the amount that catalyzed the release of 1 μmol of *p*-nitrophenol per min from *p*NP-β-GalNAc under experimental conditions. Values represent the mean of technical triplicate measurements. For Assay II (4MU-β-GalNAc as a substrate), the reaction mixture was formulated using 10 nmol of 4MU-β-GalNAc and an appropriate amount of enzyme in 100 μL of a 100 mM buffer solution. Following incubation for 30 min, the fluorescence intensity was measured using a Synergy 2 multimode microplate reader (BioTek) at excitation and emission wavelengths of 360 and 460 nm, respectively. Values represent the mean of technical triplicate measurements. In Assay III (oligosaccharides as substrates), reaction mixtures containing 5 nmol of oligosaccharides and an appropriate amount of enzyme in 20 μL of 100 mM buffer were incubated for 16 hours. The samples were boiled for 5 min to stop the reaction. The samples were dried, and the residues were dissolved in 10 μL of a methanol:water (1:1, v/v) solution and applied to a TLC plate, which was then developed using a 1-butanol:acetic acid:water (2:1:1, v/v/v) solution. Oligosaccharides and GalNAc were visualized using a diphenylamine-aniline-phosphate reagent (a mixture of 0.4 g diphenylamine, 0.4 mL aniline, 3 mL 85% phosphoric acid, and acetone [20 mL]). The optimal pH was determined using the GTA buffer [50 mM 3,3-dimethyl glutaric acid, 50 mM tris(hydroxymethyl)aminomethane, and 50 mM 2-amino-2-methyl-1,3-propanediol]. To avoid thermal degradation of *p*NP-substrates, the optimum temperatures of the enzymes were maintained between 0 and 75℃. The substrate specificity of the enzymes was examined using the following *p*NP-glycosides (50 nmol) or 4MU-glycosides (10 nmol): GalNAc-β- or α-*p*NP, Galβ1-3GalNAc-β- or α-*p*NP, GlcNAc-β- or α-*p*NP, galactose-β- or α-*p*NP, glucose- β- or α-*p*NP, arabinose-β- or α-*p*NP, mannose-β- or α-*p*NP, fucose-β- or α-*p*NP, xylose-β- or α-*p*NP, sulfate-*p*NP, GalNAc-β-4MU, GalNAc4S-β-4MU and GalNAc6S-β-4MU.

### Protein thermal shift assay

The protein thermal shift assay was conducted in the StepOnePlus Real-Time PCR System (Applied Biosystems) using Applied Biosystems Protein Thermal Shift Dye. The *T*_m_ of the proteins was calculated using the Protein Thermal Shift software v1.4 (Applied Biosystems).

### Crystallization, data collection, and structure determination

Purified β-NGA was concentrated to 10–20 mg/mL. This sample solution (0.5 μL) was mixed with an equivalent volume of the reservoir solution. Crystallization was performed by the sitting drop vapor diffusion technique at 20°C. Crystals were first formed in the crystallization screening trial and were reproduced by seeding the crystals into solutions prepared in an identical manner (Supplementary Data S7). For crystallization of the β-NGA-(GalNAc-thiazoline) complex, GalNAc-thiazoline was incorporated into the β-NGA protein solution to obtain a final concentration of 5 mM. GalNAc-thiazoline was prepared as previously described (31). A crystallization solution containing 20% (v/v) glycerol was used as a cryoprotectant for X-ray diffraction data collection. The X-ray diffraction experiments were performed at the BL32XU beamline of SPring-8. All diffraction data were collected using the automated data collection system ZOO (50). The obtained data were processed with XDS (51) using the automated data processing pipeline KAMO (52). For the X-ray diffraction data of NgaDssm complexed with GalNAc-thiazoline, automated structural analysis was performed using NABE (Matsuura et al., under review), and the structure was solved by molecular replacement using the AlphaFold2 model (48). PHENIX (53), COOT (54), and REFMAC (55) were employed for structure refinement. Molecular images were displayed using PyMol (Schrödinger LLC, Palo Alto, CA, USA). The secondary structural elements in Supplementary Figs. 1–5 were determined using the ESPript software (56).

## Author Contributions

1. T. Sumida conceived and designed the study; performed molecular experiments, protein purifications, enzyme characterization, and protein crystallization; determined the protein structures; and wrote the manuscript. S.H. performed the bioinformatics analyses and wrote the manuscript. K.U. performed the molecular experiments and protein purifications. A.I. and K.T. performed NMR analysis. T. Sengoku performed structural prediction. K.A.S. synthesized inhibitors. S.D. and T.N. wrote the manuscript and supervised the project. S.F. determined the protein structures and wrote the manuscript. All authors reviewed the manuscript draft and approved the final manuscript.

## Acknowledgements

We would like to express sincere appreciation to the captain, crew, and all onboard scientists and technicians of the KR11-11 cruise. We are extremely grateful to the ROV *ABISMO* development and operation teams. Computations were partially performed on the NIG supercomputer at the ROIS National Institute of Genetics and the Data Analysis System and the Earth Simulator at JAMSTEC. This research was funded by the Research Support Project for Life Science and Drug Discovery (Basis for Supporting Innovative Drug Discovery and Life Science Research (BINDS)) from AMED under Grant Number JP22ama121001 (*support number 3118*). The synchrotron radiation experiments were performed at the BL32XU of SPring-8 with the approval of the Japan Synchrotron Radiation Research Institute (JASRI) (Proposal No. 2021A6700). We thank Kunio Hirata and BL32XU beamline staff for assisting with X-ray crystallographic data collection and analysis. We are grateful to Naohiro Matsugaki for the diffraction data analysis, Yoshitaka Moriwaki for the AlphaFold2 analysis, and Yukishige Ito for the NMR analysis. We thank Yasuhiro Shimane, Miho Hirai, and Fumie Kondo for their assistance in this study. This work was supported by the Japan Society for the Promotion of Science (Grant Numbers JP20K15444, JP22K05398) and the Mizutani Foundation for Glycoscience grant. K.A.S. appreciates the support of the Australian Research Council (FT100100291).

## Data availability

The metagenomic sequencing data were deposited in the DDBJ Sequence Read Archive under BioProject ID PRJDB15058 [https://ddbj.nig.ac.jp/resource/bioproject/PRJDB15058]. All coordinates are deposited in the PDB under accession numbers 8K2F, 8K2G, 8K2H, 8K2I, 8K2J, 8K2K, 8K2L, 8K2M, and 8K2N. All the requisite data for evaluating the conclusions are present in the article and/or the Materials and Methods. Additional information related to this study may be requested from the authors.

---

**Extended Data Fig. 1.**
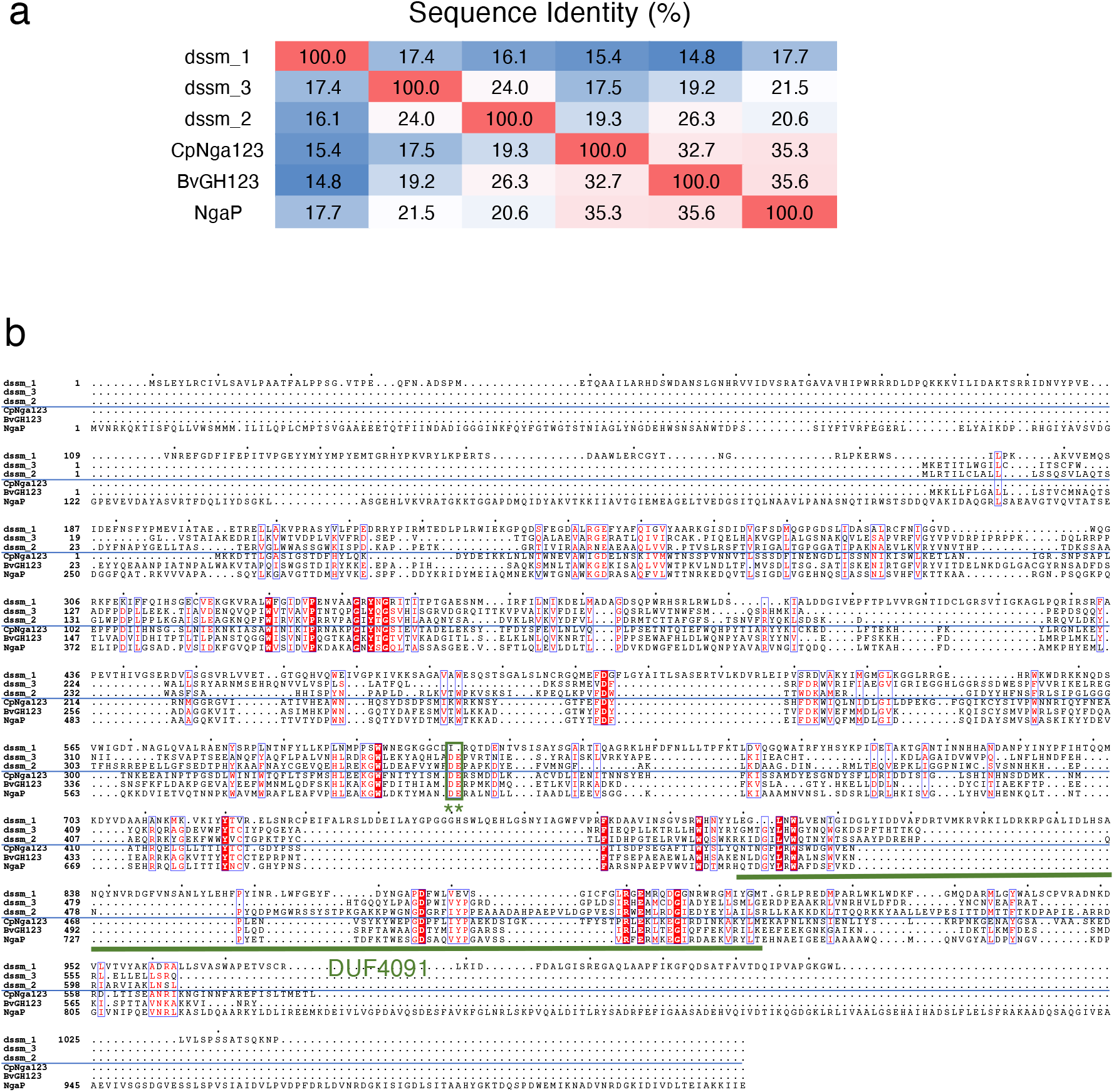
Candidate β-NGA sequences retrieved from deep-sea sediment metagenomes. **a**, Sequence identities within β-NGA candidates retrieved from deep-sea sediment metagenomes and the known GH123 genes. **b**, Alignment of gene sequence. Residues conserved in all the proteins are shown on a red background. A conserved DE motif (green asterisk, *) is indicated by a green box. The DUF4091 is underlined.

**Extended Data Fig. 2.**
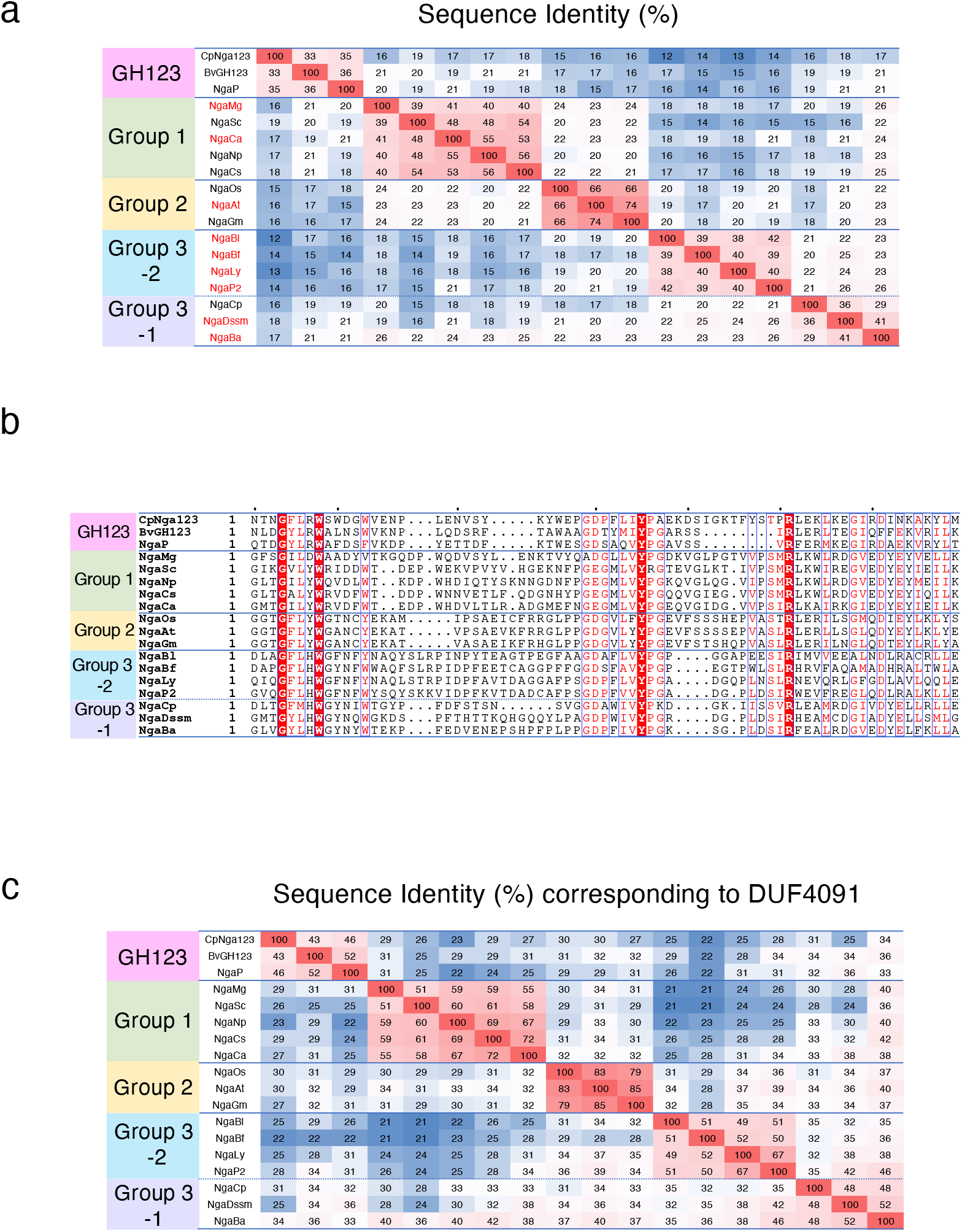
Candidate β-NGA sequences containing DUF4091. **a**, Sequence identities within β-NGA genes. Names in red represent the enzymes whose activities were experimentally confirmed in this study. **b**, Alignment of the gene sequences corresponding to DUF4091. **c**, Sequence identities corresponding to DUF4091 with known GH123 and β-NGA candidates.

**Extended Data Fig. 3.**
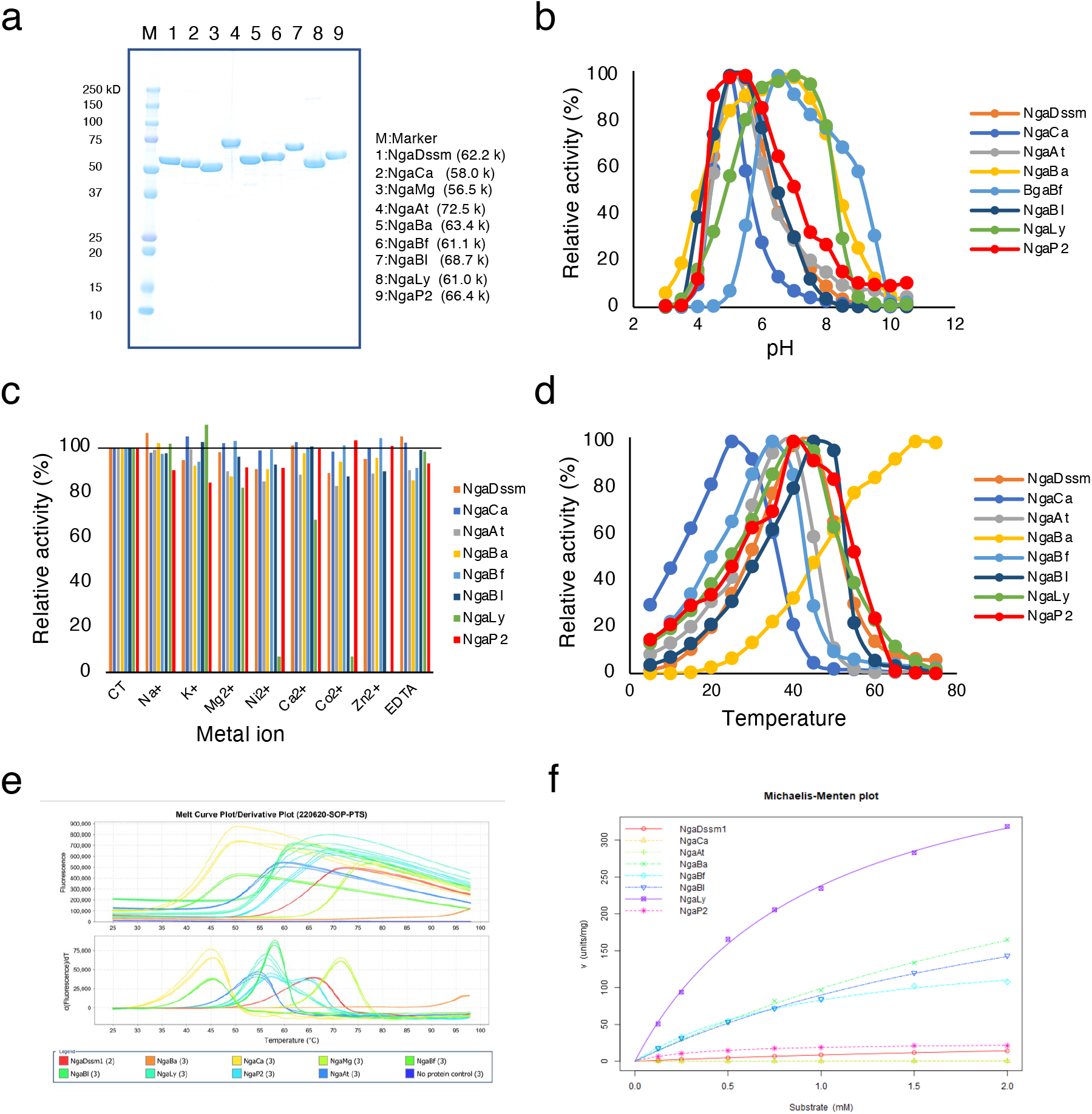
General properties of recombinant β-NGAs. **a**, SDS-PAGE analysis of recombinant β-NGAs, stained with Coomassie brilliant blue. Effect of **b**, pH and **c**, metal ions on enzymatic activity of recombinant β-NGAs at optimal temperature. **d**, Effect of temperature on enzymatic activity of recombinant β-NGAs. **e**, Protein thermal shift data of recombinant β-NGAs. **f**, s-v plots of recombinant β-NGAs. All values represent the mean of triplicate measurements.

**Extended Data Fig. 4.**
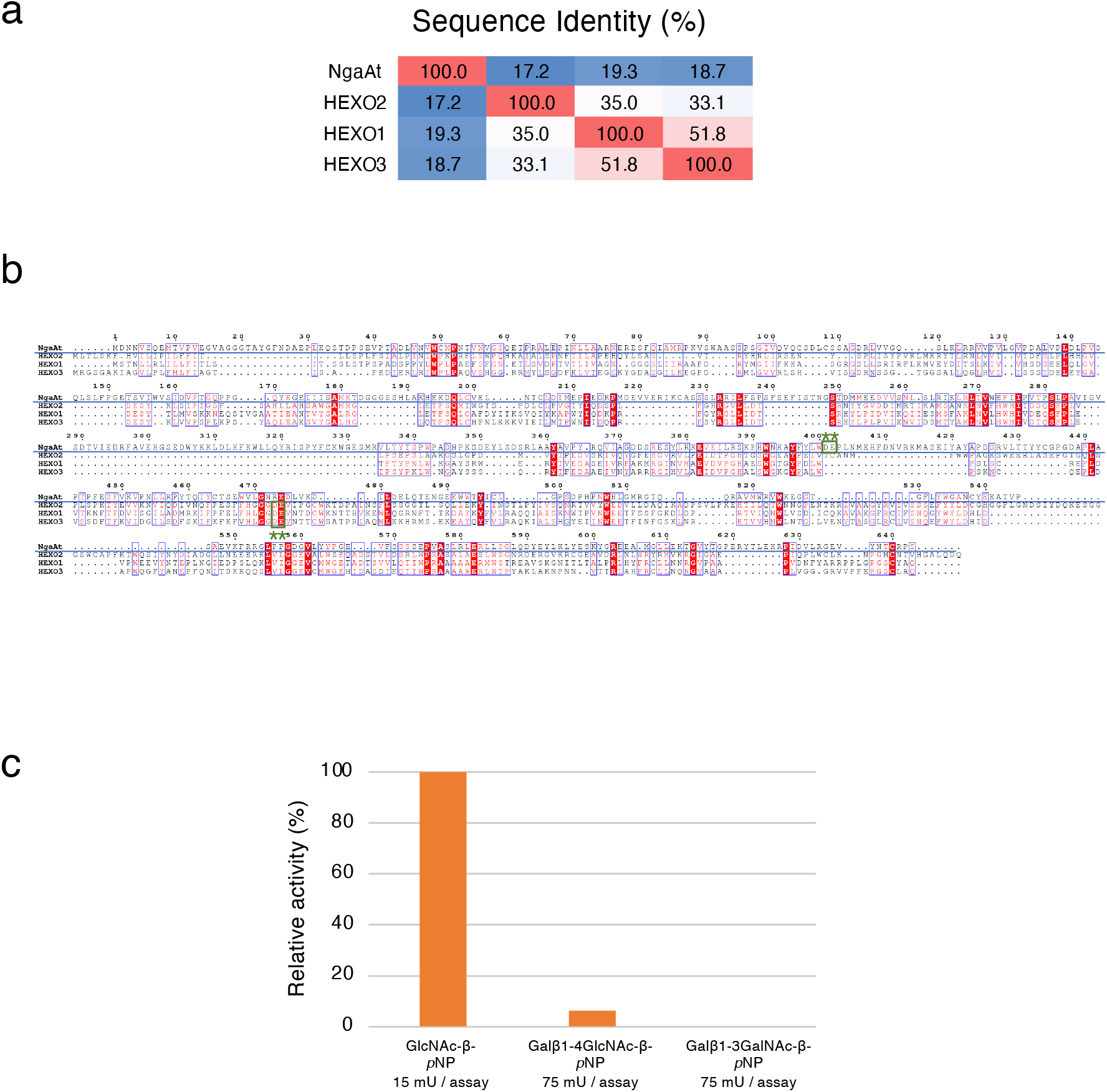
Comparison between NgaAt and HEXO1–3. **a**, Sequence identities within the NgaAt and HEXO1–3 genes. **b**, Alignment of gene sequences. Residues conserved in all the proteins are displayed in a red background. The DE motifs (green asterisk, *) of NgaAt and HEXO1–3 are indicated by green boxes. **c**, Specificity of NgaAt against different *p*NP-substrates. Values represent the mean of technical triplicate measurements.

**Extended Data Fig. 5.**
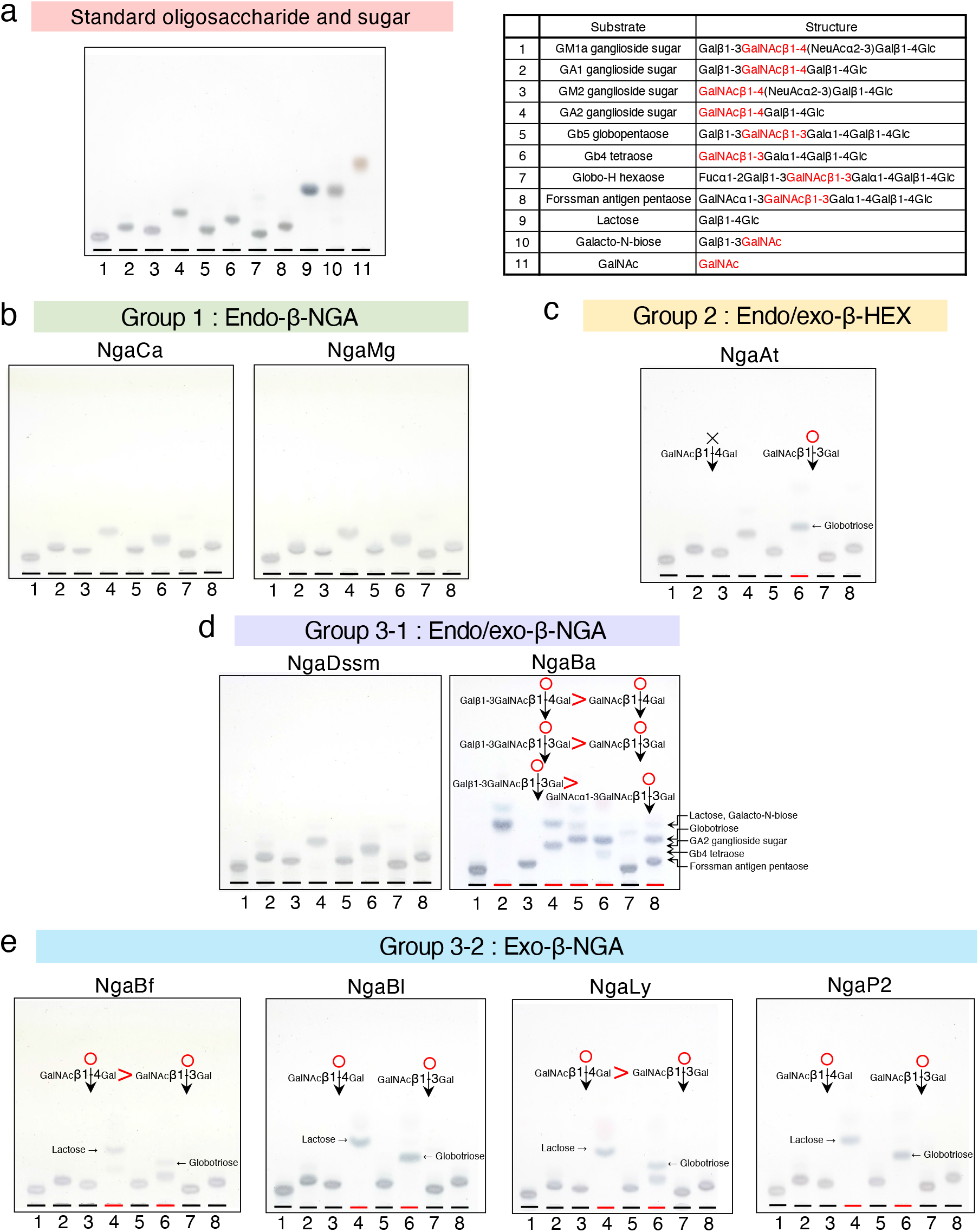
Hydrolysis of oligosaccharides by recombinant β-NGAs. **a**, TLC analysis of the standard oligosaccharides and sugars (left), together with the structures of the oligosaccharides used in the TLC assay (right). **b–e**, TLC demonstrates the hydrolysis of oligosaccharides of GM1a, asialo GM1 (GA1), GM2, asialo GM2 (GA2), Gb5, Gb4, Globo H, and Forssmann antigens. Oligosaccharides in each lane were arranged in the same order as in the standard TLC.

**Extended Data Fig. 6.**
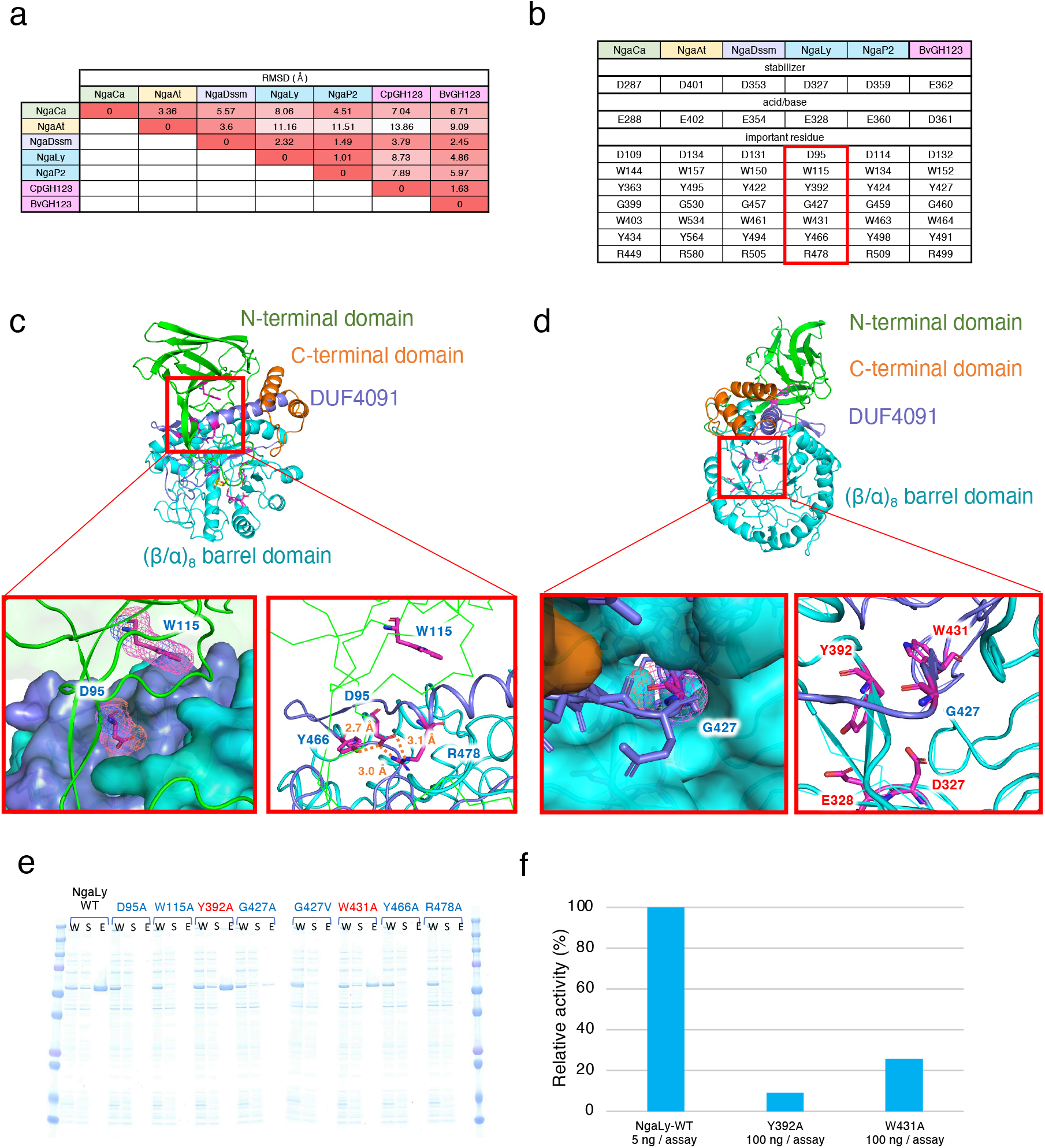
Comparison of overall structure and effect of the mutation on amino acids conserved throughout all β-NGA groups. **a**, Structural similarity between β-NGA groups compared using the root mean square distance (RMSD). **b**, Amino acids conserved between β-NGA groups. Point mutation experiments were performed on the amino acids of NgaLy as specified in the red box. **c**, **d**, Detailed views of interactions between conserved residues (**c**, D95, W115, Y466, R478, **d**, G427). The β-sandwich domain at the N-terminus, the (β/α)_8_-barrel domain, DUF4091, and the C-terminal domain are depicted in green, cyan, purple, and orange, respectively. Magenta sticks represent the conserved amino acids. The ligands are indicated by yellow sticks. **e**, SDS-PAGE analysis of point mutants (W: whole-cell lysate; S: supernatant protein after sonication and centrifugation; E: eluted protein after affinity purification). The amino acids that likely contribute to structural stability and those involved in substrate recognition are demarcated with blue and red colors, respectively. **f**, Relative activity of the NgaLy Y392A and W431A mutants compared to that of the wild-type (WT) enzyme. Values represent the mean of technical triplicate measurements.

**Extended Data Fig. 7.**
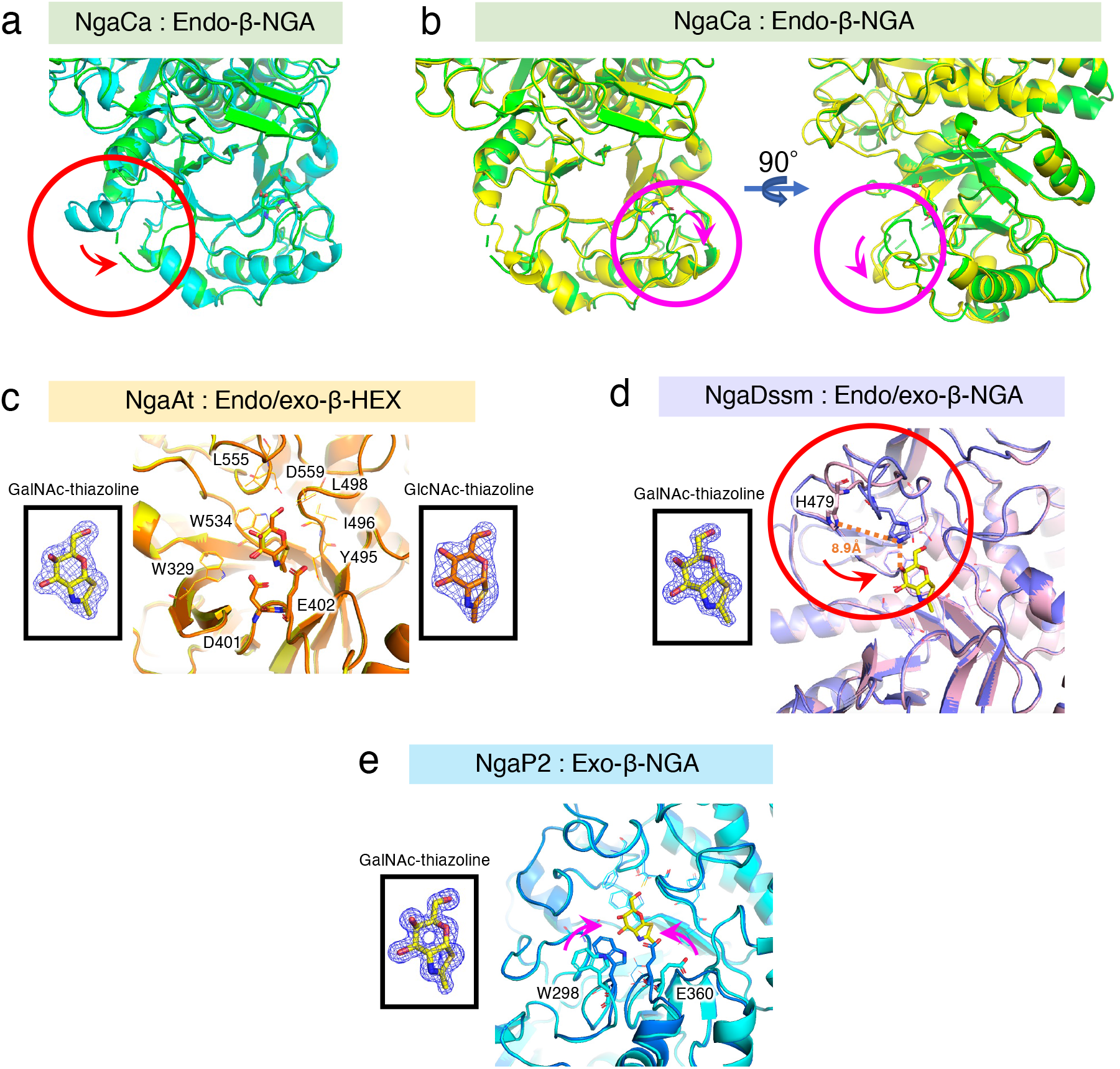
The structures of NgaCa, NgaAt, NgaDssm, and NgaP2. **a**, Comparison between the apo1 crystal structure (green) and AlphaFold2 predicted structure (cyan) of NgaCa. The position of the loop behind the second β-sheet in the (β/α)_8_-barrel domain is demarcated with a red circle. The DE motif is indicated by a stick. **b**, Distinction between apo 1 (green) and apo 2 (yellow) in NgaCa. The apo 2 structure is derived from crystals to which 5 mM Galβ1-3GalNAc was incorporated during crystallization, but Galβ1-3GalNAc is not visible. However, the loops surrounding the active site are different from those surrounding apo 1, and the loops move as the active site cleft expands (magenta circle). These two distinct states were designated as closed (apo 1) and open (apo 2), respectively. The DE motif is indicated by a stick. **c**, Superimposition of the GalNAc- and GlcNAc-thiazoline bond forms of NgaAt (yellow and orange, respectively). The positions of the amino acids involved in substrate recognition are identical. GalNAc- and GlcNAc-thiazoline are represented by yellow and orange sticks, respectively. Polder maps of GalNAc- and GlcNAc-thiazoline (4σ) are illustrated as blue meshes. **d**, Superimposition of the apo (pink) and the GalNAc-thiazoline-bound forms (purple) of NgaDssm. In the apo form, His479 is distant from the active site. In the GalNAc-thiazoline bond, His 479 was located within hydrogen-bonding distance from the 4-OH of GalNAc-thiazoline. These two distinct forms are designated as closed and open states, respectively. GalNAc-thiazoline is depicted by a yellow stick. A polder map of GalNAc-thiazoline (4σ) is displayed as a blue mesh. **e**, Superimposition of the apo form (cyan) of NgaP2 onto its GalNAc-thiazoline-bound form (blue). The residues that shifted upon GalNAc-thiazoline binding, Trp298 and Glu360, are depicted as sticks. The displacements of these residues are indicated by arrows. GalNAc-thiazoline is represented by a yellow stick. A polder map of GalNAc-thiazoline (4σ) is illustrated as a blue mesh.

**Extended Data Fig. 8.**
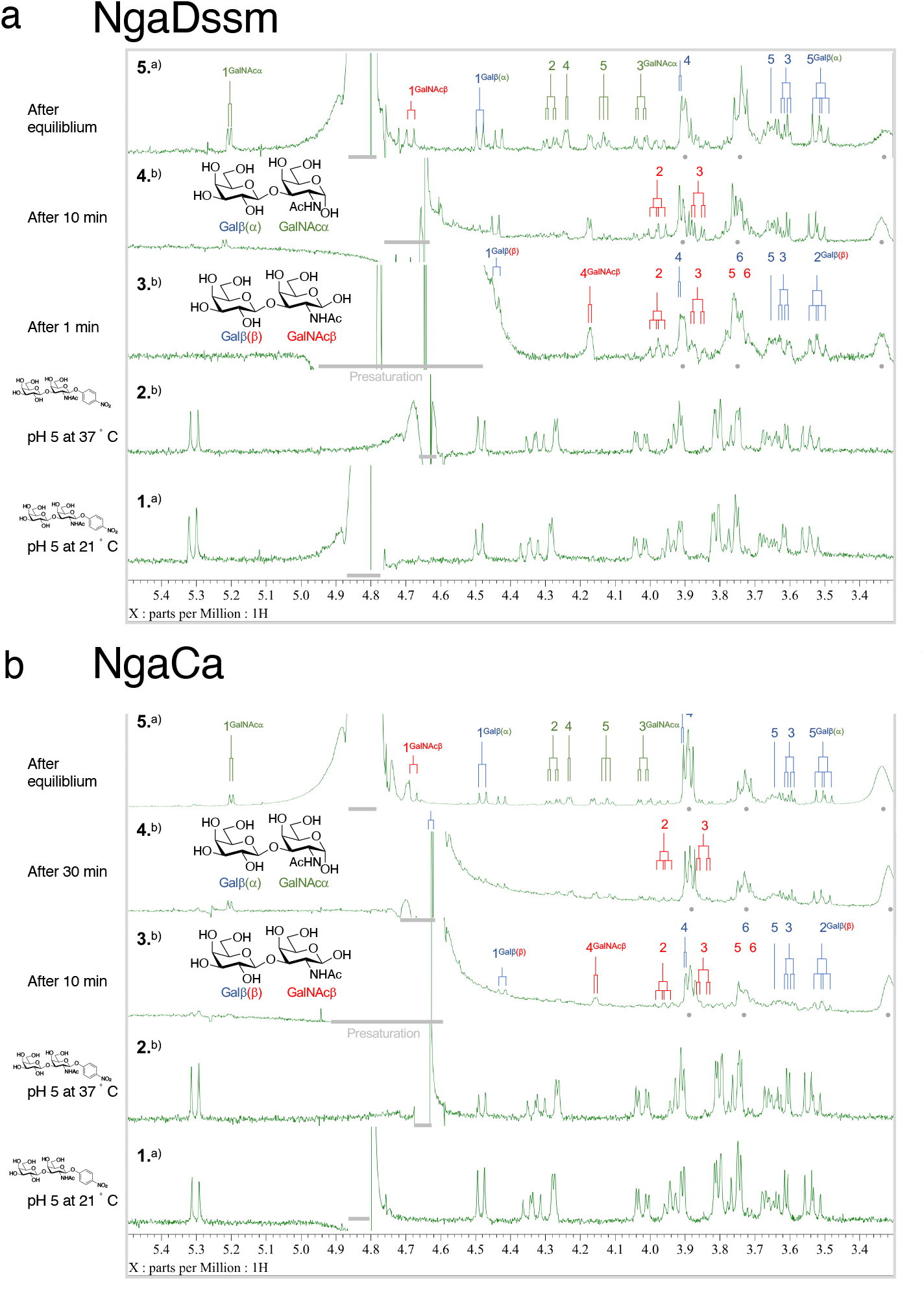
NgaDssm and NgaCa are anomer-retaining enzymes. **a**, ^1^H NMR spectrum monitoring the activity of NgaDssm toward Galβ1-3GalNAc-β-*p*NP in D_2_O/H_2_O (32:1, 100 mM citrate buffer, pH(D) 5.0) at 37℃. 1: Substrate at 21℃; 2: Substrate at 37℃ at pH(D) 5.0 before addition of enzyme solution; 3: Reaction after 1 min; 4: Reaction after 10 min; 5: Reaction after attaining equilibrium. The enzyme was dissolved in 20 mM HEPES-Na (pH 7.5), 150 mM NaCl, and 1 mM DTT in H_2_O and premixed with D_2_O (D_2_O/H_2_O = 6:1) before treatment with the substrate. **b**, ^1^H NMR spectrum monitoring the activity of NgaCa toward Galβ1-3GalNAc-β-*p*NP in D_2_O/H_2_O (11:1, 100 mM citrate buffer, pH(D) 5.0) at 37℃. 1: Substrate at 21℃; 2: Substrate at 37℃ D_2_O/H_2_O (11:1, 100 mM citrate buffer, pH(D) 5.0); 3: Reaction after 10 min; 4: Reaction after 30 min; 5: Reaction after attaining equilibrium. The enzyme was dissolved in 20 mM HEPES-Na (pH 7.5), 150 mM NaCl, and 1 mM DTT in H_2_O (32.5 mg/mL) and premixed with D_2_O (D_2_O/H_2_O = 2:1) before treatment with the substrate. a) Measured at 21℃ with presaturation at 4.80 ppm. b) Measured at 37℃ with presaturation at 4.63 ppm. The gray dots symbolize the peaks of the reagents in solution (DTT and HEPES). The gray bars indicate the areas affected by presaturation to reduce the HOD peak.

**Extended Data Fig. 9.**
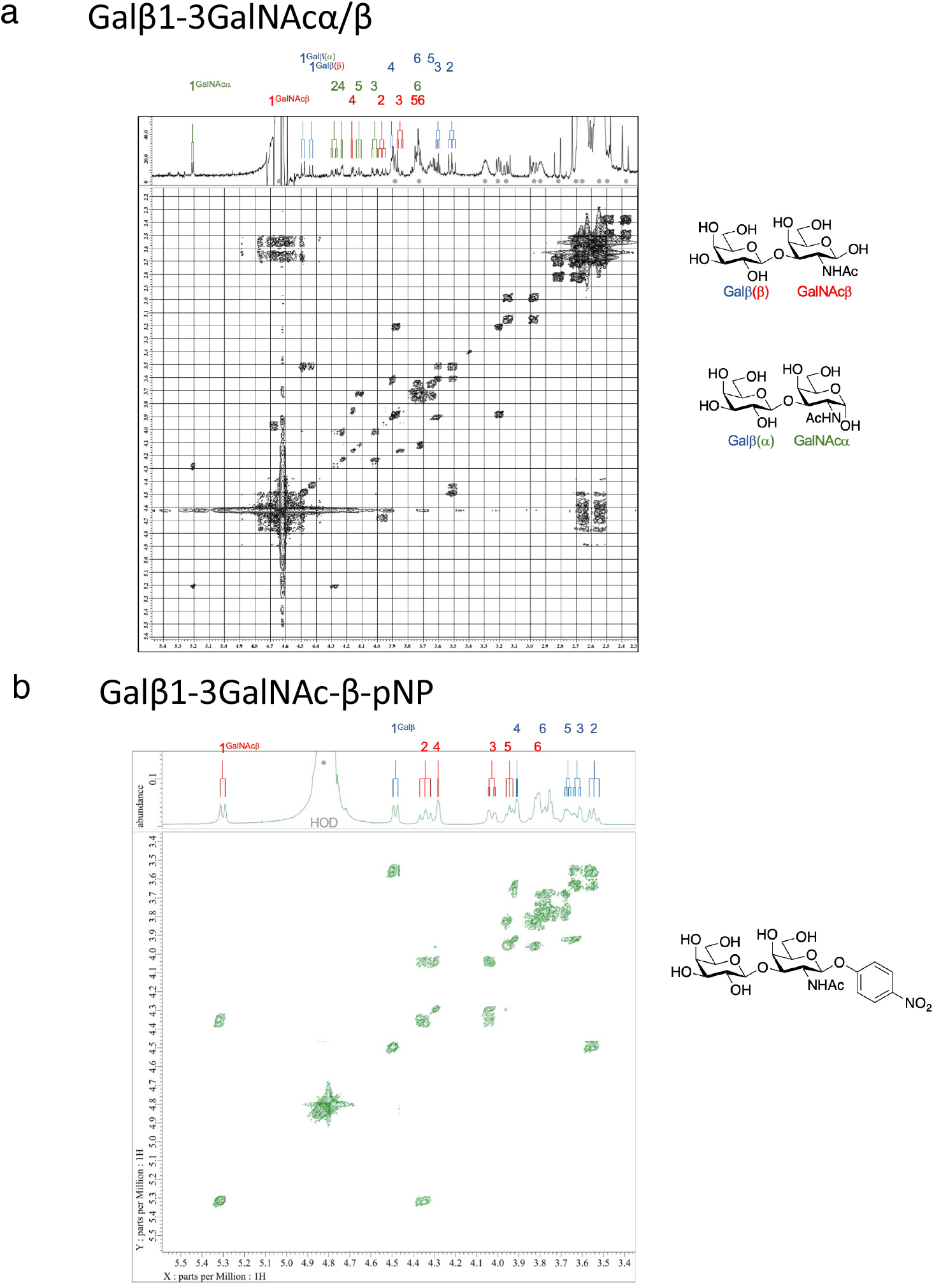
H-H COSY and ^1^H NMR spectra of Galβ1-3GalNAcα/β and Galβ1-3GalNAc-β-*p*NP. **a**, 2D H-H COSY and 1D ^1^H NMR spectra of the reaction mixtures at equilibrium after hydrolysis. The mixture comprises Galβ1-3GalNAcα/β (∼1.5:1). Assignments were carried out by 1D ^1^H NMR and 2D H-H COSY as the chemical shift of ^1^H at 1 of β-GalNAc (4.58 ppm) is overlapped under the large HOD peak (4.4–4.8). α and β in parentheses indicate the stereochemistry of the reducing GalNAc residue of the corresponding disaccharides. **b**, H-H COSY and 1D ^1^H NMR spectra of Galβ1-3GalNAc-β-*p*NP in D_2_O. Assignments were performed using 1D ^1^H and ^13^C, 2D H-H COSY, TOCSY, HMQC, HMQC-TOCSY, and HMBC.

**Extended Data Fig. 10.**
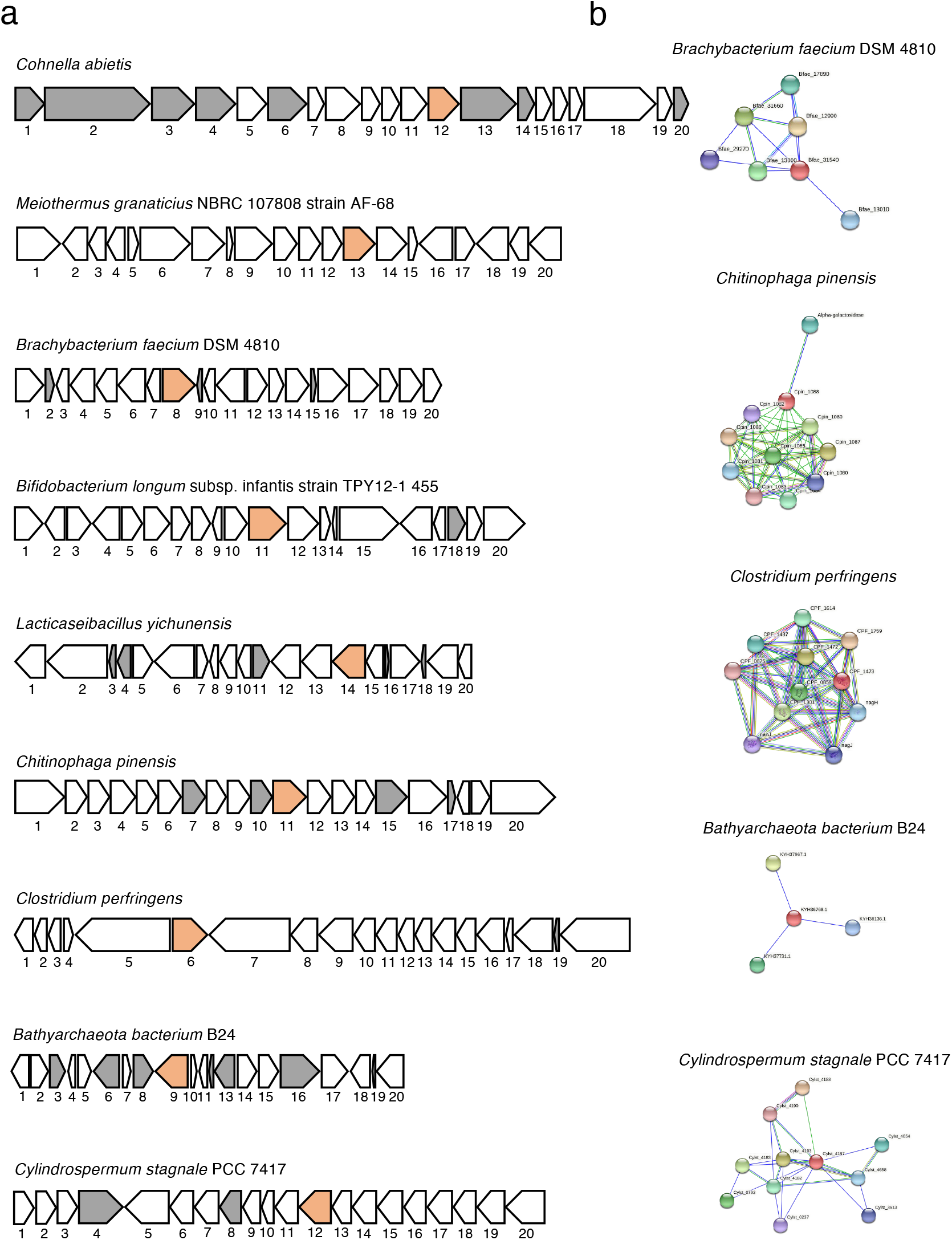
Neighborhood genes and potential protein interaction networks of β-NGAs. **a**, Neighborhood genes around the β-NGA. Functional annotations were obtained from the NCBI GenBank database, as listed in Supplementary Data S4. The β-NGAs and hypothetical proteins are depicted in orange and gray, respectively. **b**, Interaction networks of β-NGA retrieved from the STRING database (v11.5). Predicted functional partner proteins are listed in Supplementary Data S5.

