## Supplementary Figure 1-8, Supplementary Table 1-7 for "Genetic and functional diversity of β-*N*-acetylgalactosamine residue-targeting glycosidases expanded by deep-sea metagenome"

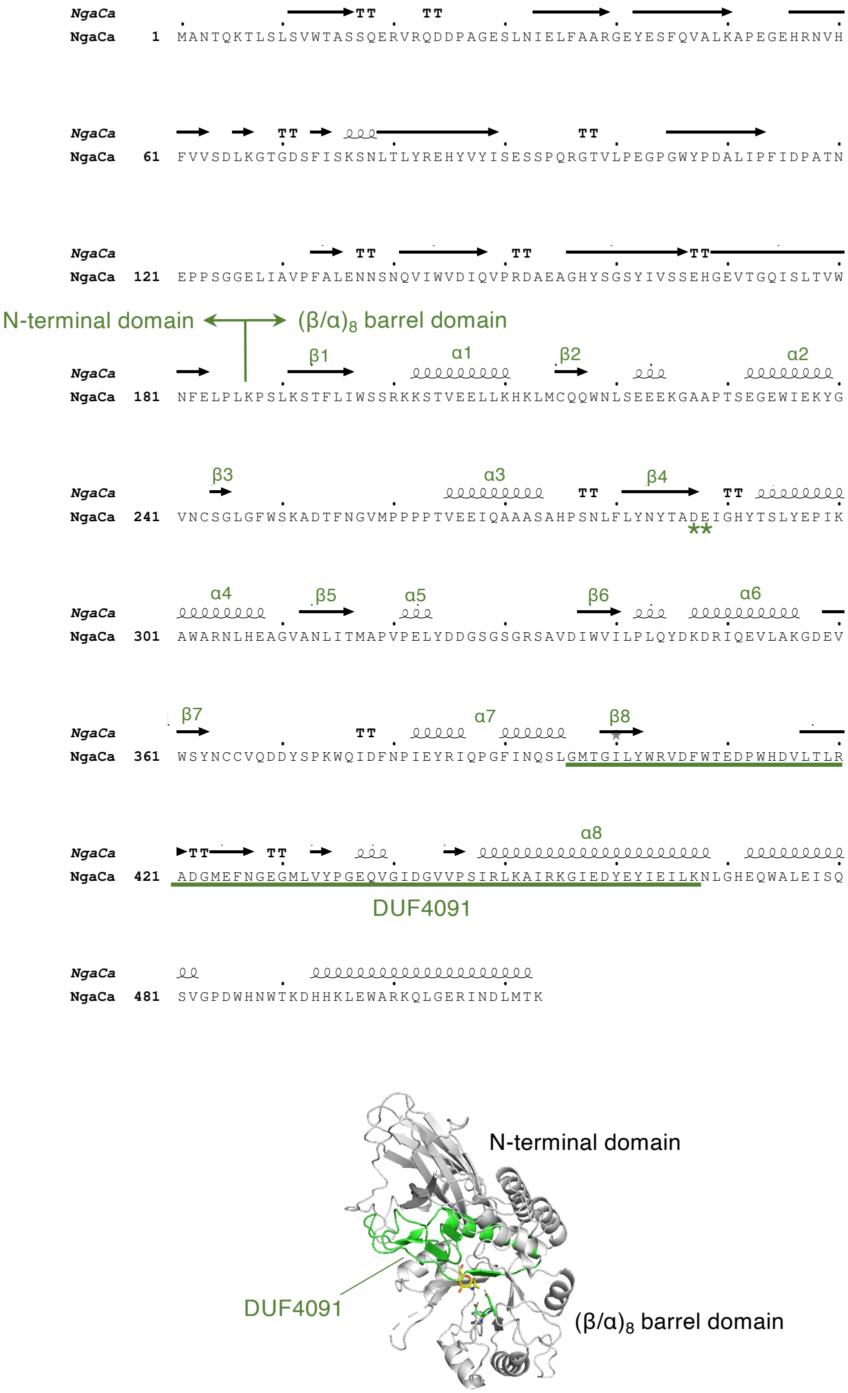


**Supplementary Figure 1. Secondary structure and characteristic domains of NgaCa (Group 1).**

The secondary structural elements were indicated above the sequence. The $(\beta/\alpha)_8$-barrel region is indicated above the secondary structural elements. The conserved DE residues (*) are indicated below the sequence. The DUF4091 region is underlined.

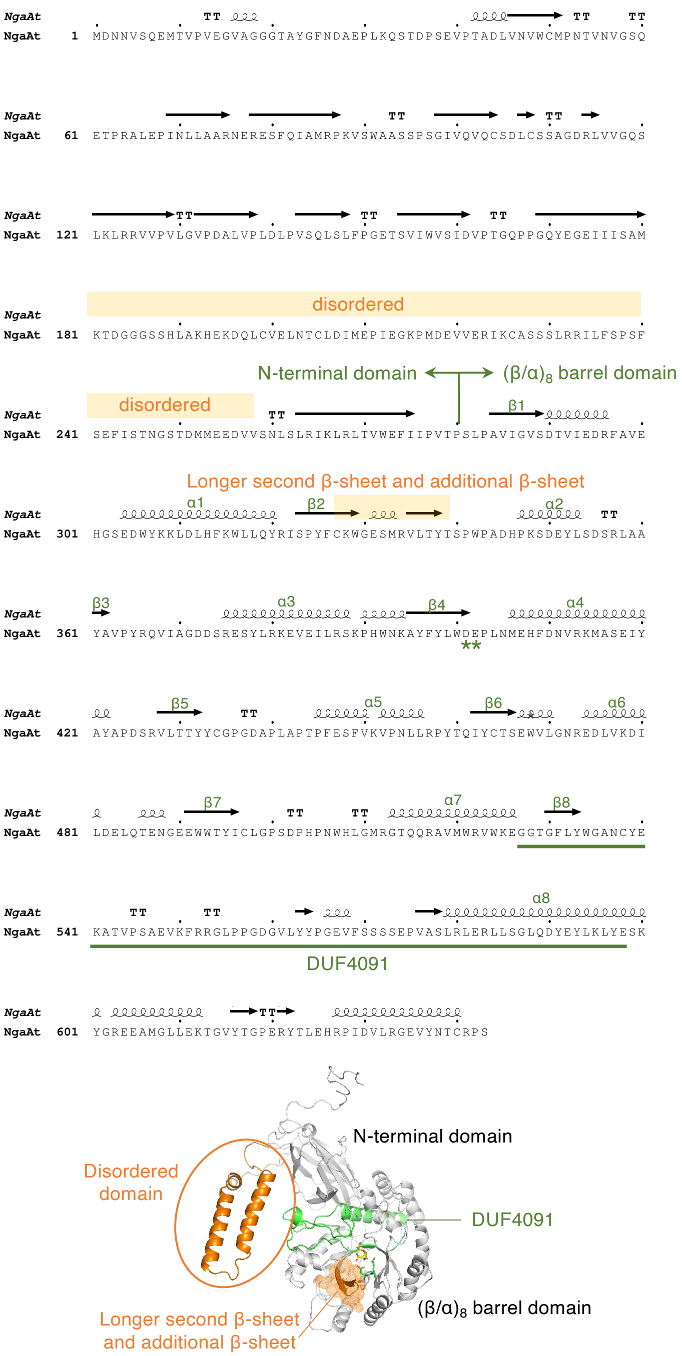


**Supplementary Figure 2. Secondary structure and characteristic domains of NgaAt (Group 2).**

The secondary structural elements were indicated above the sequence. The additional domain characteristic of Group 2 is shown in orange. The $(\beta/\alpha)_8$-barrel region is indicated above the secondary structural elements. The conserved DE residues (*) are indicated below the sequence. The DUF4091 region is underlined.

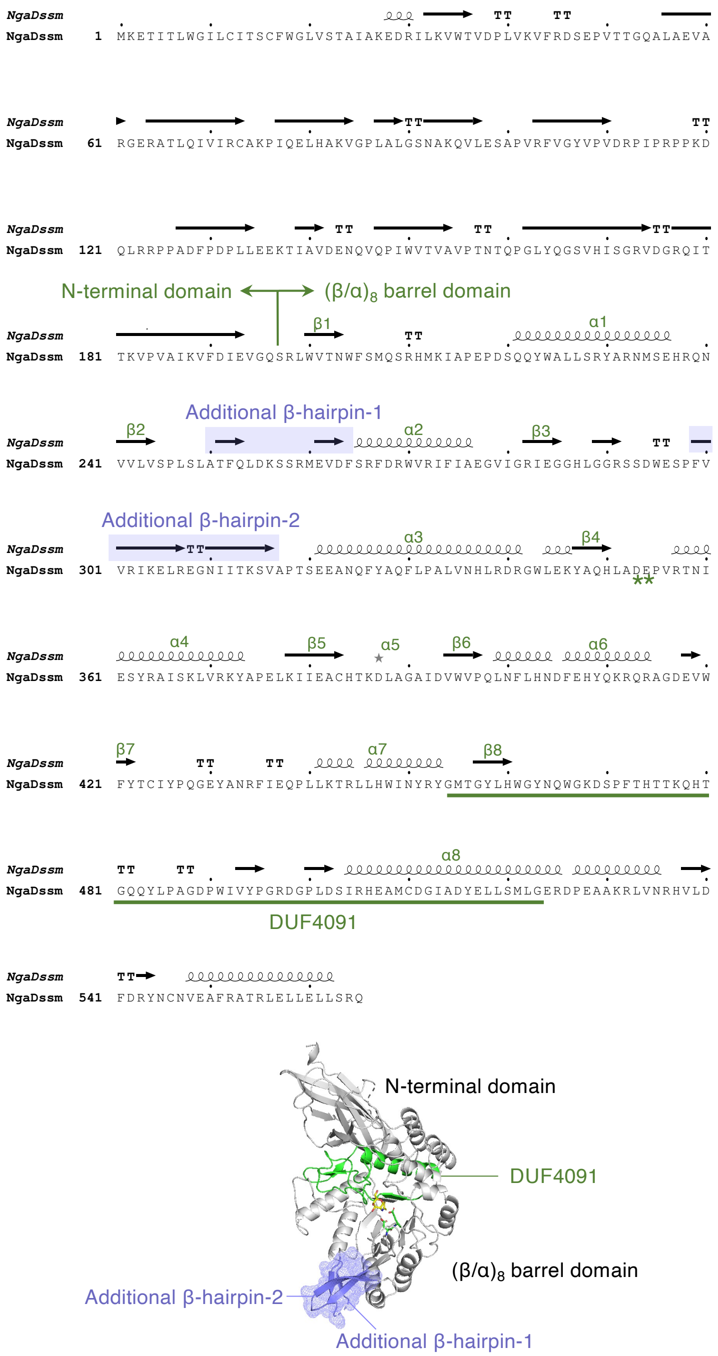


**Supplementary Figure 3. Secondary structure and characteristic domains of NgaDssm (Group 3-1).**

The secondary structural elements were indicated above the sequence. The additional domain characteristic of Group 3-1 is shown in purple. The $(\beta/\alpha)_8$-barrel region is indicated above the secondary structural elements. The conserved DE residues (*) are indicated below the sequence. The DUF4091 region is underlined.

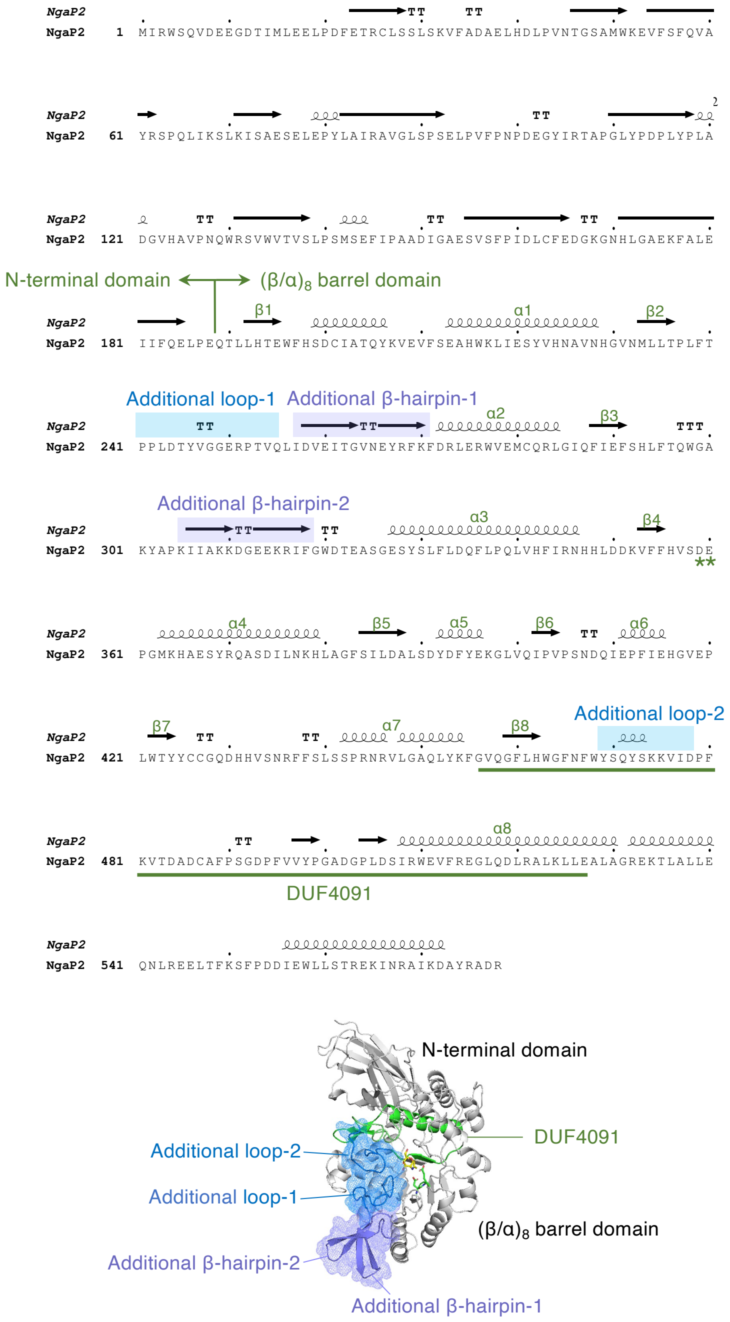


**Supplementary Figure 4. Secondary structure and characteristic domains of NgaP2 (Group 3-2).**

The secondary structural elements were indicated above the sequence. The additional β-hairpins and loops characteristic of Group 3-2 are shown in purple and blue, respectively. The $(\beta/\alpha)_8$-barrel region is indicated above the secondary structural elements. The conserved DE residues (*) are indicated below the sequence. The DUF4091 region is underlined.

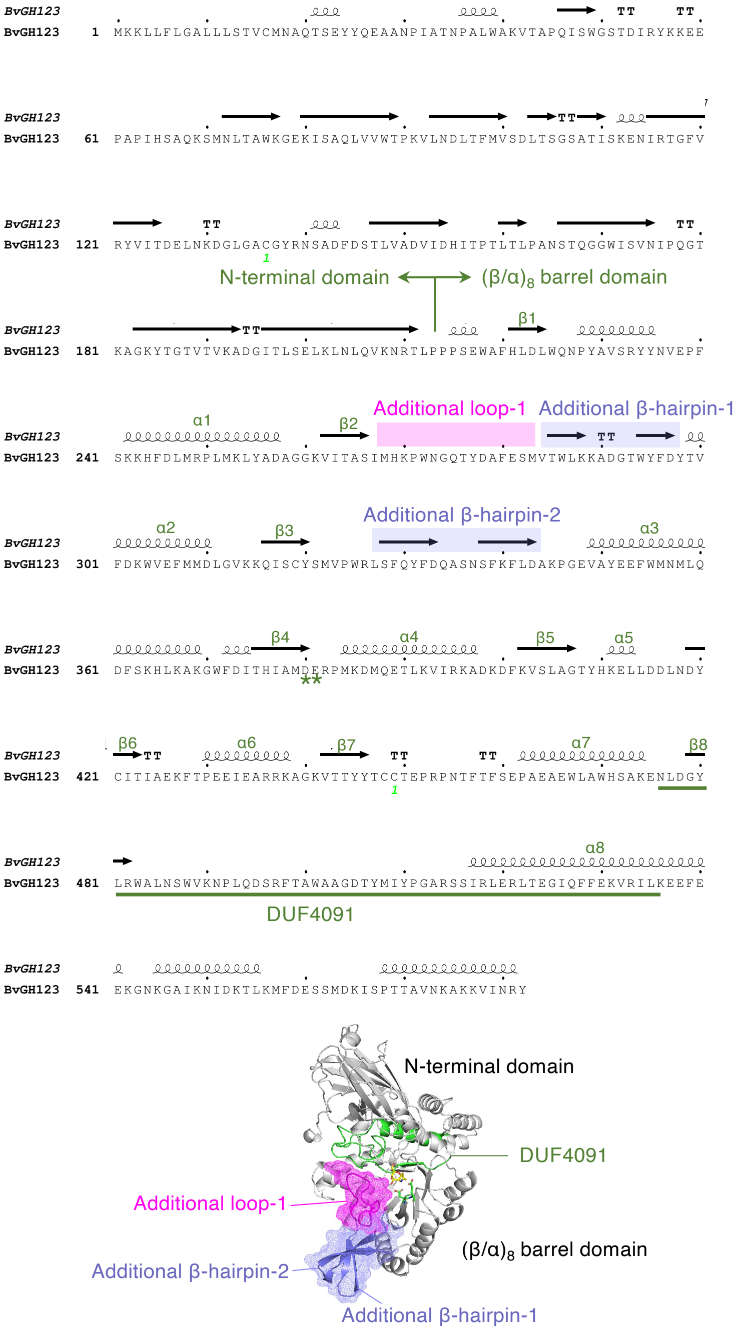


**Supplementary Figure 5. Secondary structure and characteristic domains of BvGH123 (GH123).**

The secondary structural elements were indicated above the sequence. The additional β-hairpins and loops characteristic of GH123 are shown in purple and magenta, respectively. The $(\beta/\alpha)_8$-barrel region is indicated above the secondary structural elements. The conserved DE residues (*) are indicated below the sequence. The DUF4091 region is underlined.

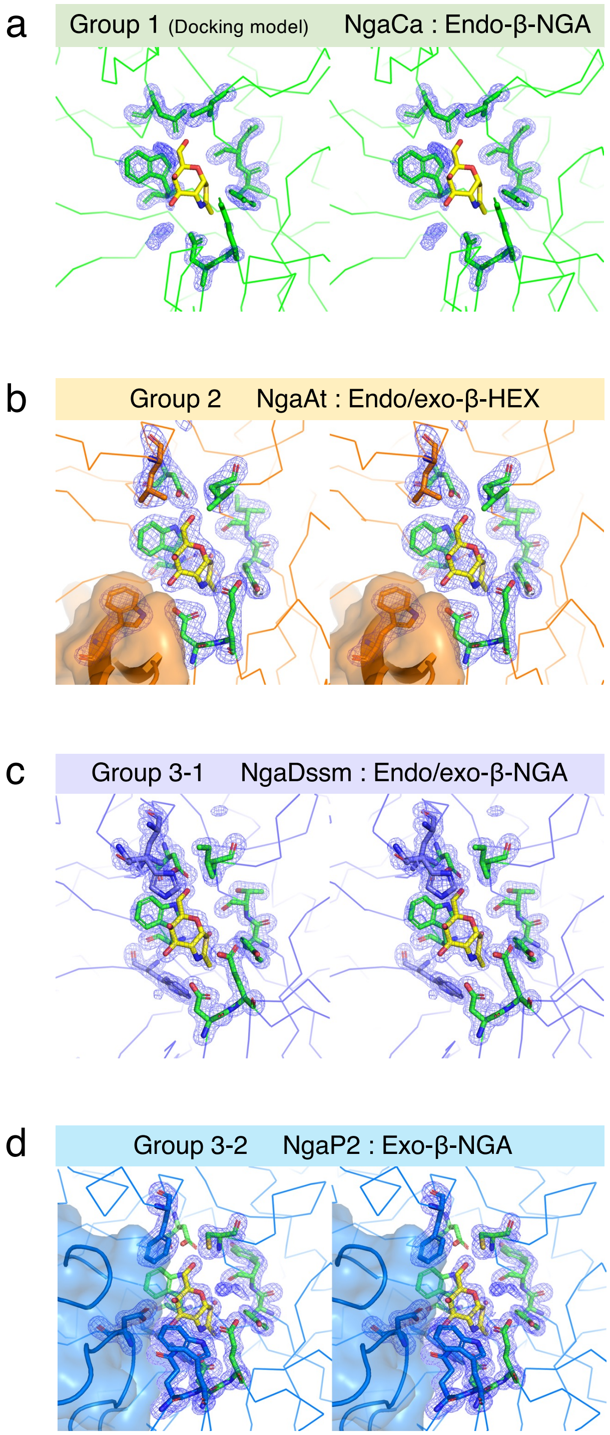


**Supplementary Figure 6 Stereo view of the active site of β-NGAs.**

**a**, A docking model of NgaCa with GalNAc-thiazoline. **b–d**, Crystal structures of each enzyme complexed with GalNAc-thiazoline. Amino acid residues and GalNAc-thiazoline are shown in stick. Polder maps (4σ) are shown as blue mesh.

**a**

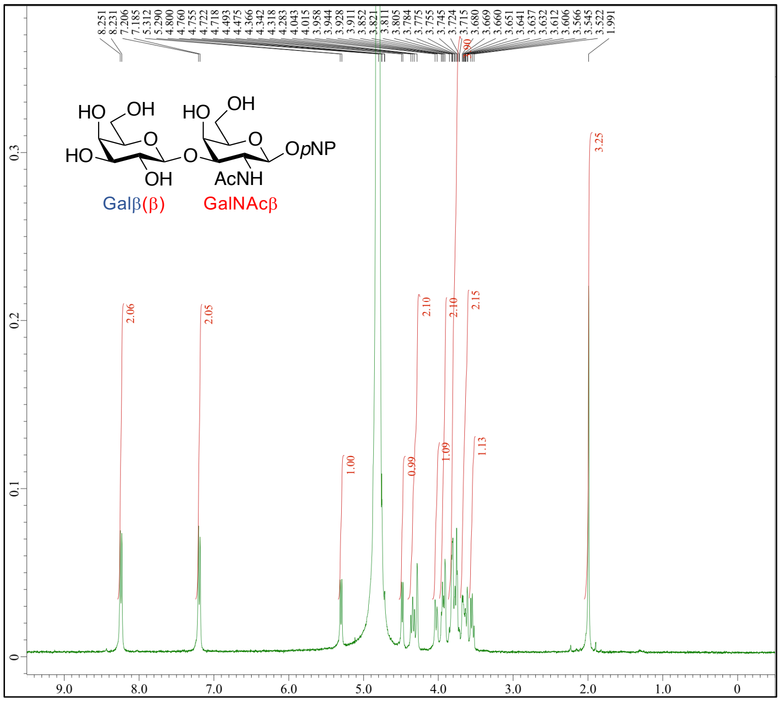


**b**

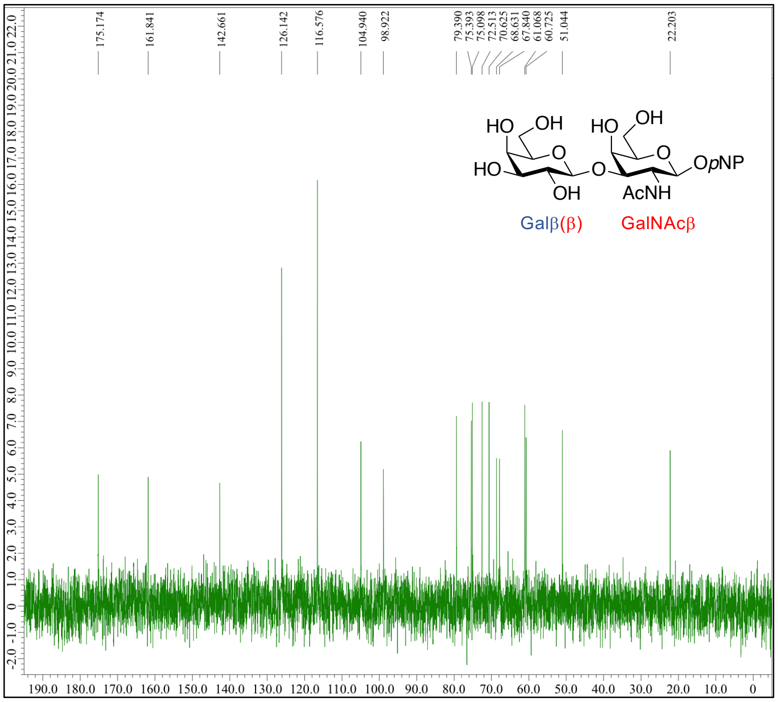


**Supplementary Figure 7. NMR spectra of Galβ1-3GalNAc-β-*p*NP in $D_2O$.**

**a**, $^{1}H$ NMR spectrum, **b**, $^{13}C$ NMR spectrum.

**c**

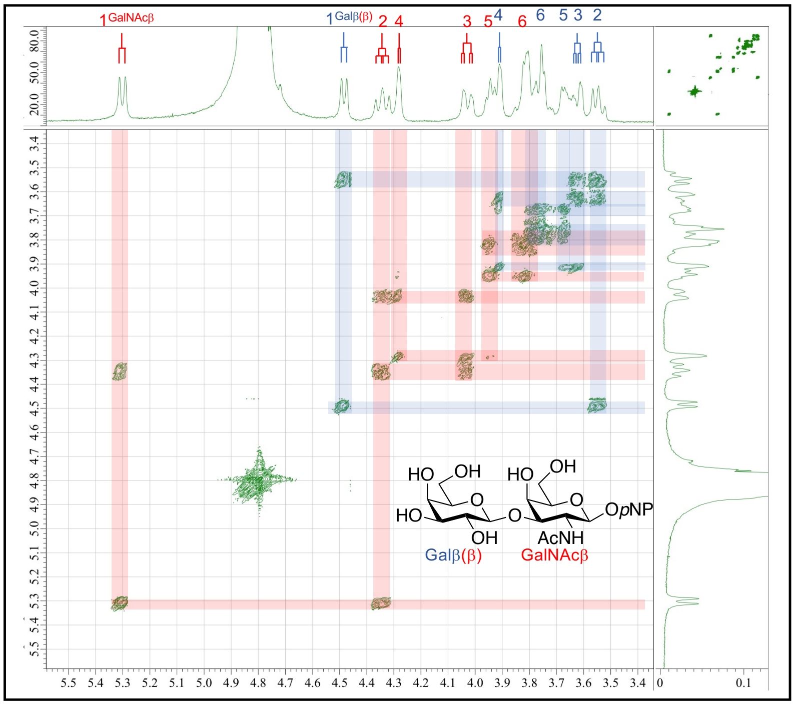


**d**

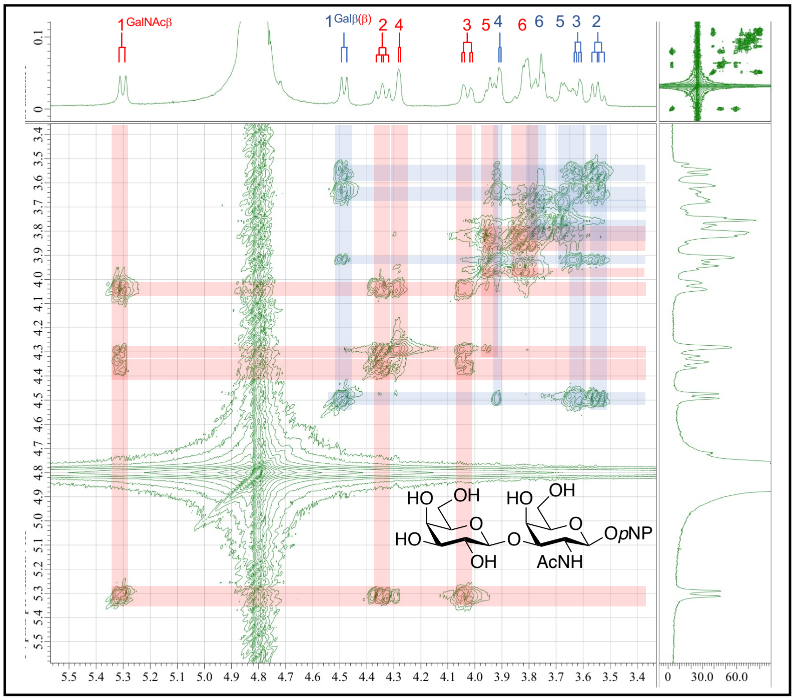


**Supplementary Figure 7. NMR spectrra of Galβ1-3GalNAc-β-*p*NP in $D_2O$.**

**c**, $^1$H–$^1$H COSY spectrum, **d**, TOCSY spectrum. Assignments are shown with the multiplications and the position numbers in each residue on the top.

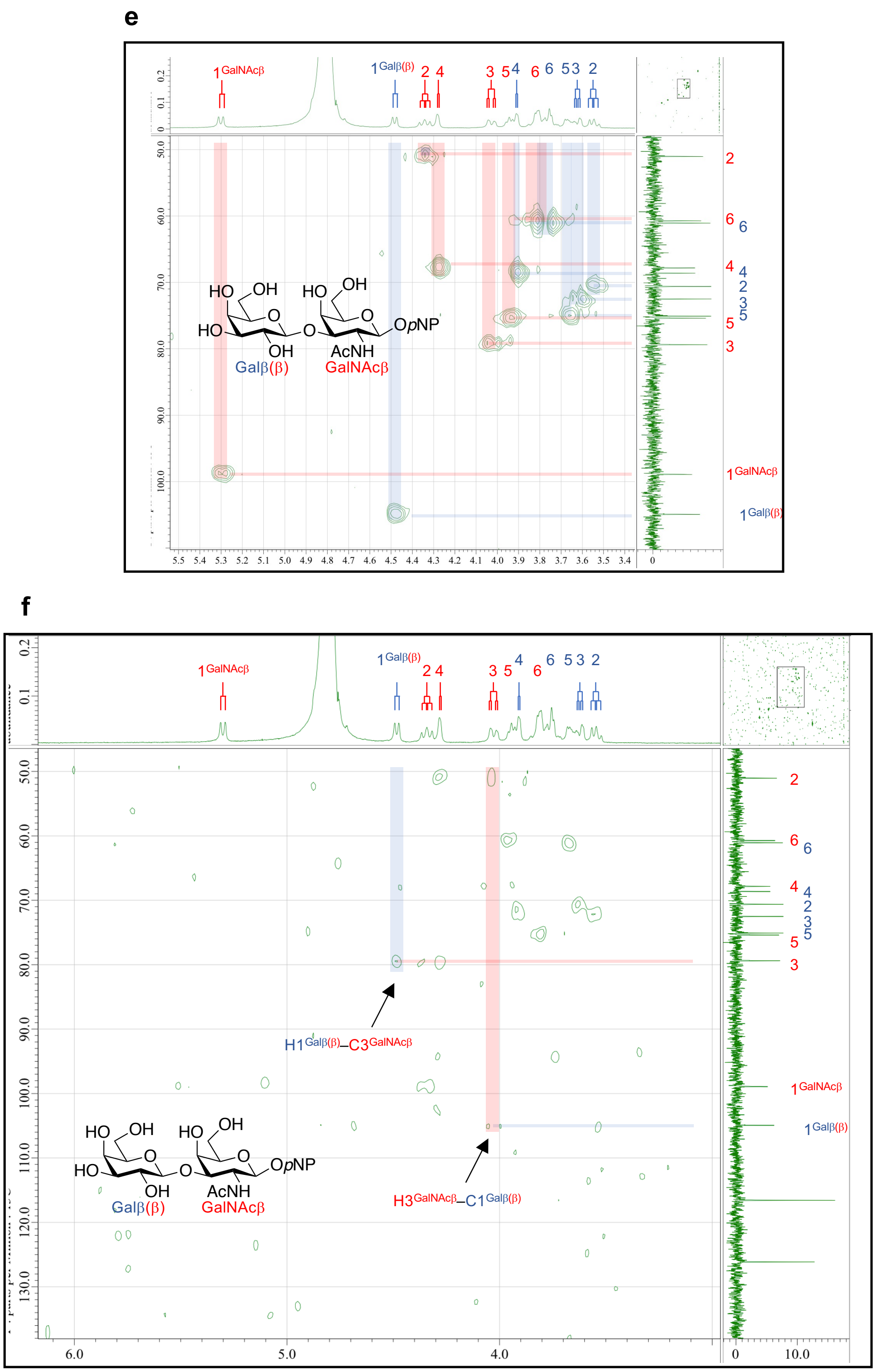


**Supplementary Figure 7. NMR spectrra of Galβ1-3GalNAc-β-*p*NP in $D_2O$.**

**e**, HMQC spectrum, **f**, HMBC spectrum. Assignments are shown with the multiplications for $^{1}$H NMR and the position numbers for both $^{1}$H and $^{13}$C NMRs in each residue.

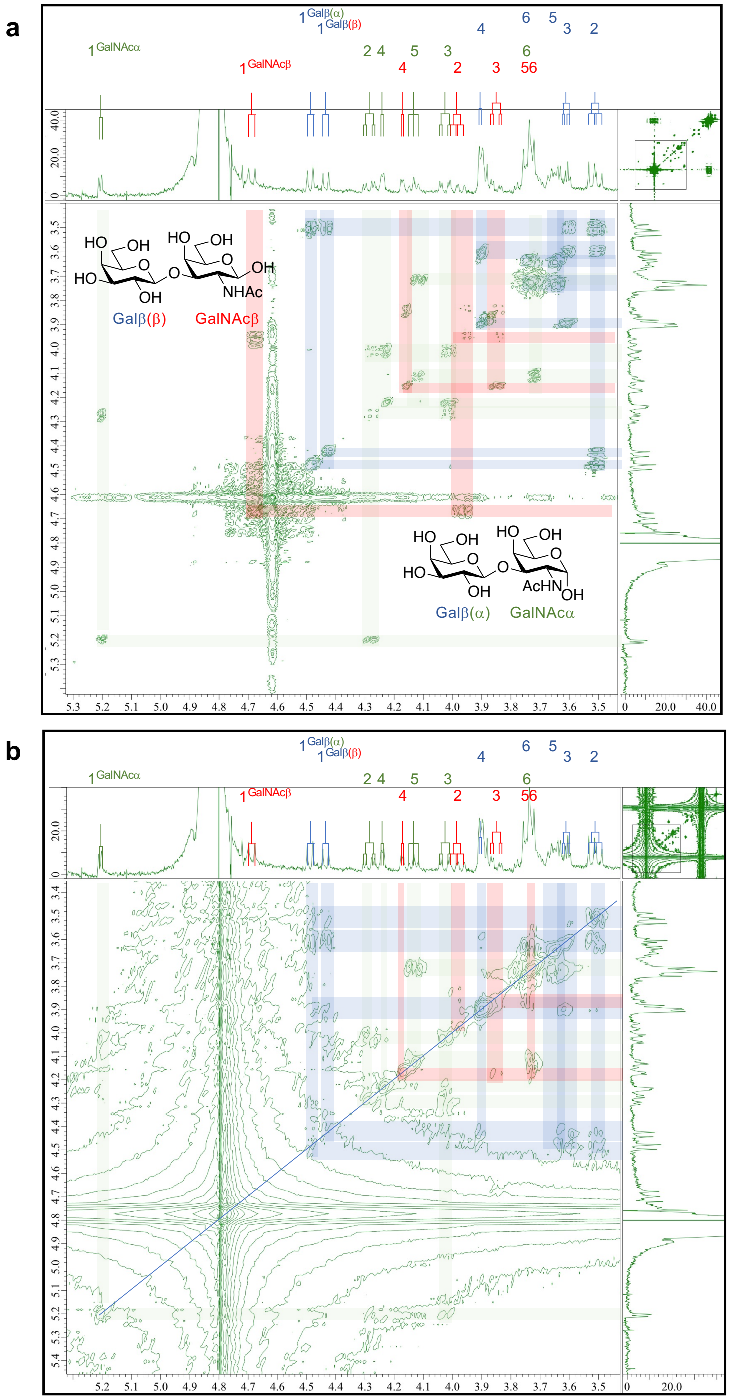


**Supplementary Figure 8. NMR spectra of Galβ1-3GalNAc-β-*p*NP treated with the enzyme at pH 5 in $D_2O$.** **a**, $^1$H–$^1$H COSY spectrum, **b**, TOCSY spectrum. The reaction mixtures after 30 min and 24 h were used for the experiments. For $^1$H–$^1$H COSY spectrum the same $^1$H NMR spectra of the reaction mixture after 24 h was used. Assignments are shown with the multiplications and the position numbers of each residue on the top.

c

d

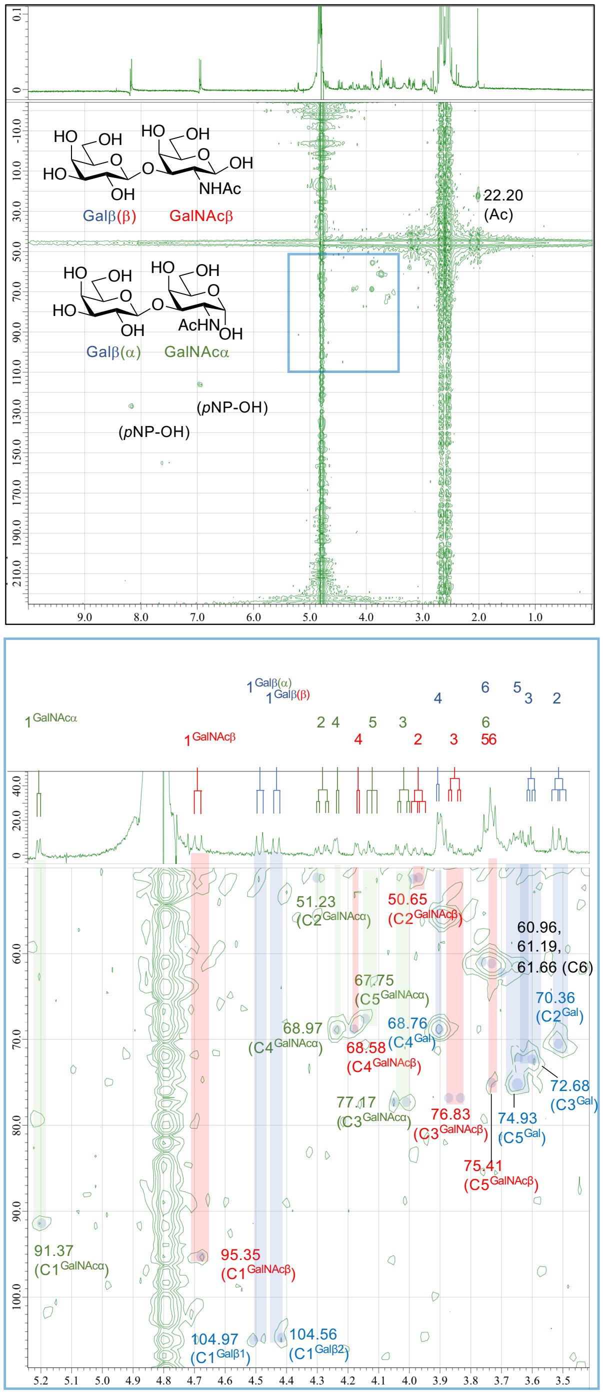


**Supplementary Figure 8. NMR spectrra of Galβ1-3GalNAc-β-*p*NP treated with the enzyme at pH in $D_2O$.**

**c**, HMQC spectrum, **d**, $^{13}$C assignment using HMQC spectrum. The reaction mixture after 24 h was used for the experiments. Assignments are shown with the multiplications for $^{1}$H NMR and the position numbers for both $^{1}$H and $^{13}$C NMRs of each residue.

**Supplementary Table 1**. Data collection and refinement statistics of the crystallography of NgaCa (Group 1).

| **Data set** | **NgaCa apo 1** | **NgaCa apo 2 (GNB soaked)** |
|---|---|---|
| **Data collection** | | |
| Beamline | BL32XU | BL32XU |
| Wavelength (Å) | 1.0000 | 1.0000 |
| Space group | $P2_1$ | $P1$ |
| Unit cell (Å/°) | $a$ = 48.699, $b$ = 70.614, $c$ = 71.604, β = 100.856 | $a$ = 53.214, $b$ = 69.679, $c$ = 71.113, α = 88.413, β = 74.329, γ = 87.925 |
| Resolution (Å) | 49.83–1.75 (1.78–1.75) | 48.36–1.75 (1.78–1.75) |
| $R_{merge}$ | 0.122 (0.511) | 0.093 (0.327) |
| $R_{pim}$ | 0.049 (0.204) | 0.067 (0.228) |
| Total reflections | 338,363 (18,019) | 341,066 (14,776) |
| Unique reflections | 47,285 (2,505) | 96,240 (4,648) |
| Mean $I/\sigma(I)$ | 10.4 (4.7) | 7.1 (3.7) |
| $CC_{1/2}$ | 0.993 (0.942) | 0.963 (0.872) |
| Completeness (%) | 98.4 (95.4) | 97.1 (93.6) |
| Multiplicity | 7.2 (7.2) | 3.5 (3.4) |
| Molecules/asymmetric unit | 1 | 2 |
| **Refinement** | | |
| Resolution (Å) | 49.88–1.75 | 47.67–1.75 |
| No. of reflections (all/free) | 47,231/2,396 | 96,224/4,810 |
| $R_{work}/R_{free}$ | 0.174/0.215 | 0.156/0.193 |
| Number of atoms | 4,222 | 8,766 |
| RMSD from ideal values | | |
| Bond lengths (Å) | 0.0102 | 0.0103 |
| Bond angles (°) | 1.65 | 1.70 |
| Clashscore | 2.71 | 2.03 |
| Molprobity score | 1.46 | 1.21 |
| Ramachandran plot (%) | | |
| Favored/allowed/outlier | 96.34/2.85/0.81 | 96.92/2.29/0.80 |
| PDB code | 8K2F | 8K2G |

Values in parentheses are for highest resolution shell.

**Supplementary Table 2**. Data collection and refinement statistics of the crystallography of NgaAt (Group 2).

| Data set | NgaAt GalNAc-thiazoline | NgaAt GlcNAc-thiazoline |
|---|---|---|
| **Data collection** | | |
| Beamline | BL32XU | BL32XU |
| Wavelength (Å) | 1.0000 | 1.0000 |
| Space group | $P2_12_12_1$ | $P2_12_12_1$ |
| Unit cell (Å/°) | *a* = 68.852, *b* = 134.036, *c* = 149.969 | *a* = 69.284, *b* = 133.833, *c* = 150.723 |
| Resolution (Å) | 48.02–2.20 (2.25–2.20) | 48.13–2.50 (2.58–2.50) |
| $R_{merge}$ | 0.226 (3.004) | 0.270 (3.022) |
| $R_{pim}$ | 0.064 (0.873) | 0.105 (1.158) |
| Total reflections | 972,501 (60,305) | 685,113 (64,372) |
| Unique reflections | 71,289 (4,507) | 49,358 (4,468) |
| Mean $I/\sigma(I)$ | 9.5 (1.9) | 7.4 (1.4) |
| $CC_{1/2}$ | 0.997 (0.648) | 0.994 (0.684) |
| Completeness (%) | 100.0 (100.0) | 100.0 (100.0) |
| Multiplicity | 13.6 (13.4) | 13.9 (14.4) |
| Molecules/asymmetric unit | 2 | 2 |
| **Refinement** | | |
| Resolution (Å) | 48.07–2.20 | 48.13–2.50 |
| No. of reflections (all/free) | 71,198/3,690 | 49,278/2,489 |
| $R_{work}/R_{free}$ | 0.169/0.229 | 0.157/0.230 |
| Number of atoms | 9,285 | 9,217 |
| RMSD from ideal values | | |
| Bond lengths (Å) | 0.0015 | 0.0149 |
| Bond angles (°) | 2.14 | 2.21 |
| Clashscore | 3.88 | 4.96 |
| Molprobity score | 1.73 | 2.02 |
| Ramachandran plot (%) | | |
| Favored/allowed/outlier | 97.04/2.24/0.72 | 96.40/3.06/0.54 |
| PDB code | 8K2H | 8K2I |

Values in parentheses are for highest resolution shell.

**Supplementary Table 3**. Data collection and refinement statistics of the crystallography of NgaDssm (Group 3-1).

| Data set | NgaDssm apo | NgaDssm GalNAc-thiazoline |
|---|---|---|
| **Data collection** [a] | | |
| Beamline | BL32XU | BL32XU |
| Wavelength (Å) | 1.0000 | 1.0000 |
| Space group | $P4_32_12$ | $P4_32_12$ |
| Unit cell (Å/°) | $a = b = 53.379$, $c = 427.352$ | $a = b = 53.537$, $c = 428.292$ |
| Resolution (Å) | 47.75–1.75 (1.78–1.75) | 47.88–1.75 (1.78–1.75) |
| $R_{merge}$ | 0.202 (1.872) | 0.286 (3.351) |
| $R_{pim}$ | 0.043 (0.410) | 0.061 (0.740) |
| Total reflections | 1,489,931 (75,841) | 1,515,006 (79,728) |
| Unique reflections | 64,770 (3,512) | 65,283 (3,545) |
| Mean $I/\sigma(I)$ | 10.6 (2.2) | 8.7 (2.3) |
| $CC_{1/2}$ | 0.997 (0.688) | 0.994 (0.629) |
| Completeness (%) | 100.0 (100.0) | 100.0 (100.0) |
| Multiplicity | 23.0 (21.6) | 23.2 (22.5) |
| Molecules/asymmetric unit | 1 | 1 |
| **Refinement** | | |
| Resolution (Å) | 47.80–1.76 | 47.93–1.75 |
| No. of reflections (all/free) | 63,071/3,242 | 65,066/3,313 |
| $R_{work}/R_{free}$ | 0.180/0.213 | 0.176/0.206 |
| Number of atoms | 4,718 | 4,307 |
| RMSD from ideal values | | |
| Bond lengths (Å) | 0.0154 | 0.0109 |
| Bond angles (°) | 2.01 | 1.65 |
| Clashscore | 3.38 | 2.90 |
| Molprobity score | 1.63 | 1.21 |
| Ramachandran plot (%) | | |
| Favored/allowed/outlier | 95.60/3.25/1.15 | 95.60/3.63/0.76 |
| PDB code | 8K2J | 8K2K |

Values in parentheses are for highest resolution shell.

**Supplementary Table 4**. Data collection and refinement statistics of the crystallography of NgaP2 (Group 3-2).

| **Data set** | **NgaP2 apo** | **NgaP2 GalNAc-thiazoline** |
|---|---|---|
| **Data collection** [a] | | |
| Beamline | BL32XU | BL32XU |
| Wavelength (Å) | 1.0000 | 1.0000 |
| Space group | *C*2 | *C*2 |
| Unit cell (Å/°) | *a* = 119.116, *b* = 60.375, *c* = 100.285, β = 120.079 | *a* = 119.975, *b* = 61.608, *c* = 100.453, β = 121.425 |
| Resolution (Å) | 46.66–1.95 (2.00–1.95) | 47.10–1.65 (1.68–1.65) |
| $R_{merge}$ | 0.137 (0.398) | 0.139 (0.565) |
| $R_{pim}$ | 0.056 (0.172) | 0.056 (0.229) |
| Total reflections | 312,050 (21,086) | 519,667 (25,463) |
| Unique reflections | 45,106 (3,144) | 74,805 (3,646) |
| Mean $I/\sigma(I)$ | 13.2 (10.3) | 8.8 (3.6) |
| $CC_{1/2}$ | 0.986 (0.541) | 0.987 (0.906) |
| Completeness (%) | 99.9 (99.6) | 99.3 (98.1) |
| Multiplicity | 6.9 (6.7) | 6.9 (7.0) |
| Molecules/asymmetric unit | 1 | 1 |
| **Refinement** | | |
| Resolution (Å) | 46.70–1.95 | 47.15–1.65 |
| No. of reflections (all/free) | 45,102/2,287 | 74,698/3,800 |
| $R_{work}/R_{free}$ | 0.148/0.195 | 0.168/0.195 |
| Number of atoms | 4,900 | 4,971 |
| RMSD from ideal values | | |
| Bond lengths (Å) | 0.0152 | 0.0120 |
| Bond angles (°) | 1.88 | 1.69 |
| Clashscore | 1.46 | 0.90 |
| Molprobity score | 1.37 | 0.88 |
| Ramachandran plot (%) | | |
| Favored/allowed/outlier | 97.31/2.15/0.54 | 97.67/1.62/0.72 |
| PDB code | 8K2L | 8K2M |

Values in parentheses are for highest resolution shell.

**Supplementary Table 5**. Data collection and refinement statistics of the crystallography of NgaLy (Group 3-2).

| Data set | NgaLy apo |
|---|---|
| **Data collection** [a] | |
| Beamline | BL32XU |
| Wavelength (Å) | 1.0000 |
| Space group | *P*1 |
| Unit cell (Å/°) | *a* = 82.930, *b* = 94.620, *c* = 116.180, α = 70.925, β = 73.805, γ = 72.429 |
| Resolution (Å) | 49.01–2.50 (2.54–2.50) |
| $R_{merge}$ | 0.099 (0.298) |
| $R_{pim}$ | – (–) |
| Total reflections | 215,567 (10,612) |
| Unique reflections | 107,859 (5,311) |
| Mean $I/\sigma(I)$ | 7.1 (2.8) |
| $CC_{1/2}$ | 0.963 (0.672) |
| Completeness (%) | 100.0 (100.0) |
| Multiplicity | 2.0 (2.0) |
| Molecules/asymmetric unit | 4 |
| **Refinement** | |
| Resolution (Å) | 48.79-2.50 |
| No. of reflections (all/free) | 107,859/5,287 |
| $R_{work}/R_{free}$ | 0.183/0.241 |
| Number of atoms | 17,554 |
| RMSD from ideal values | |
| Bond lengths (Å) | 0.00151 |
| Bond angles (°) | 2.16 |
| Clashscore | 4.61 |
| Molprobity score | 2.06 |
| Ramachandran plot (%) | |
| Favored/allowed/outlier | 95.01/4.11/0.88 |
| PDB code | 8K2N |

Values in parentheses are for highest resolution shell.

**Supplementary Table 6**. $^{1}$H and $^{13}$C NMR data of Galβ1-3GalNAcα/β.[a,b)] The Galβ1-3GalNAcα/β used for the measurement were in the reaction mixture of Galβ1-3GalNAc-β-pNP with NgaDssm in $D_2O$ [pH(D) 5.0] at 37°C. The sample contains about a 1.5:1 mixture of Galβ1-3GalNAcα/β after hydrolysis. (Supplementary Figure 7).

| Gal*p*β$^{1}$(α)[c)] | $^{1}$H / ppm | *J* / Hz | $^{13}$C/ ppm[d)] | Gal*p*β$^{2}$(β)[c)] | $^{1}$H / ppm | *J* / Hz | $^{13}$C/ ppm[d)] |
|---|---|---|---|---|---|---|---|
| 1 | 4.49 | d, 7.6 | 104.97 | 1 | 4.43 | d, 8.0 | 104.56 |
| 2 | 3.51 | dd, 10.0, 7.6 | 70.36 | 2 | 3.51 | dd, 10.0, 8.0 | 70.36 |
| 3 | 3.61 | dd, 10.0, 3.6 | 72.68 | 3 | 3.61 | dd, 10.0, 3.6 | 72.68 |
| 4 | 3,90 | d, 3.6 | 68.76 | 4 | 3,90 | d, 3.6 | 68.76 |
| 5 | 3.63–3.68 | m | 74.93 | 5 | 3.63–3.68 | m | 74.93 |
| 6 | 3.70–3.79 | m | 60.96/61.19 | 6 | 3.70–3.79 | m | 60.96/61.19 |
| | 3.70–3.79 | m | /61.66 | | 3.70–3.79 | m | /61.66 |
| GalNAc*p*α | $^{1}$H / ppm | *J* / Hz | $^{13}$C/ ppm | GalNAc*p*β | $^{1}$H / ppm | *J* / Hz | $^{13}$C/ ppm |
| 1 | 5.21 | d, 3.6 | 91.37 | 1 | 4.69 | d, 9.2 | 98.92 |
| 2 | 4.34 | dd, 11.6, 3.6 | 51.23 | 2 | 3.98 | dd, 11.2, 9.2 | 50.65 |
| 3 | 4.29 | dd, 9.6, 3,2 | 71.17 | 3 | 3.84 | dd, 11.2, 3.2 | 76.83 |
| 4 | 4.02 | d, 3.2 | 68.97 | 4 | 4.17 | d, 3.2 | 68.58 |
| 5 | 4.13 | dd 6.4, 4,8 | 67.65 | 5 | 3.70–3.79 | m | 75.41 |
| 6 | 3.70–3.79 | m | 60.96/61.19 | 6 | 3.70–3.79 | m | 60.96/61.19 |
| | 3.70–3.79 | m | /61.66 | | 3.70–3.79 | m | /61.66 |
| Ac | 2.01 | s | 22.20 | Ac | 2.01 | s | 22.20 |

a) 400 MHz at 21.2 °C in $D_2O$ presaturated and referenced using the peak of HOD at 4.80 and the native scale for $^{1}$H and $^{13}$C NMRs, respectively; b) The peaks of HEPES/ DTT were appeared in $^{1}$H NMR at δ 3.90, 3.75, 3.64, 3.23, 3.15, 2.98, 2.83–2.40 ppm; c) *α* and β in parentheses indicate the stereochemistry of the reducing GalNAc residue of the corresponding disaccharides; d) $^{13}$C NMR data of the Galβ1-3GalNAcα/β were obtained using HMQC spectrum (Supplementary Figures 7 C and D).

**Supplementary Table 7**. $^{1}$H and $^{13}$C NMR data of Galβ1-3GalNAc-β-*p*NP (Supplementary Figure 8).[a)]

| D-Gal*p*β | $^{1}$H / ppm | *J* / Hz | $^{13}$C/ ppm | Key HMBC |
|---|---|---|---|---|
| 1 | 4.48 | d, 7.2 | 104.94 | → Irr. (79.3, $C3^{\text{D-GalNAc}}$) |
| 2 | 3.54 | dd, 9.2, 7.2 | 70.62 | |
| 3 | 3.62 | dd, 9.2, 2.0 | 72.51 | |
| 4 | 3,91 | br s | 68.63 | |
| 5 | 3.64–3.69 | m | 75.10 | |
| 6 | 3.93 | dd 12.4, 7.2 | 61.07 | |
| | 3.78 | dd 12.4, 3.6 | | |
| β-D-GalNAc*p* | $^{1}$H / ppm | *J* / Hz | $^{13}$C/ ppm | HMBC |
| 1 | 5.30 | d, 8.8 | 98.92 | |
| 2 | 4.34 | dd, 9.6, 8.8 | 51.04 | |
| 3 | 4.03 | dd, 9.6, 1.6 | 79.34 | → Irr. (104.94, $C1^{\text{D-Gal}p\beta}$) |
| 4 | 4.28 | br s | 67.84 | |
| 5 | 3.92–3.96 | m | 75.39 | |
| 6 | 3.84 | dd 12.4, 6.4 | 60.72 | |
| | 3.79 | dd 12.4, 5.6 | | |
| Ac | 1.99 | s | 22.20 | |
| | - | - | 175.17 | |
| *p*NP | 8.23–8.25 | m | 126.14 | |
| | 7.18–7.21 | m | 116.58 | |
| | - | - | 161.84, 142.66 | |

a) 400 MHz at 21.2 °C in $D_2O$ referenced using the peak of HOD at 4.80 and the native scale for $^{1}$H and $^{13}$C NMRs, respectively.
